# Bryophyte spermiogenesis occurs through multimode autophagic and nonautophagic degradation

**DOI:** 10.1101/2021.08.17.456730

**Authors:** Takuya Norizuki, Naoki Minamino, Hirokazu Tsukaya, Takashi Ueda

## Abstract

Mitochondria change their morphology in response to developmental and environmental cues. During sexual reproduction, bryophytes produce spermatozoids with two mitochondria in the cell body. Although intensive morphological analyses have been conducted thus far, how this fixed number of mitochondria is realized remains unknown. Here, we investigated how mitochondria are reorganized during spermiogenesis in *Marchantia polymorpha*. We found that the mitochondrial number is reduced to one through fission followed by autophagic degradation during early spermiogenesis, and then the posterior mitochondrion arises by fission of the anterior mitochondrion. Autophagy is also responsible for the removal of other organelles, including peroxisomes, but these other organelles are removed at distinct developmental stages from mitochondrial degradation. We also found that spermiogenesis involves nonautophagic organelle degradation. Our findings highlight the dynamic reorganization of mitochondria, which is regulated distinctly from that of other organelles, and multiple degradation mechanisms operate in organelle remodeling during spermiogenesis in *M. polymorpha*.

## INTRODUCTION

The shape and distribution of mitochondria change dynamically in response to developmental or environmental cues to meet cellular energy or metabolic demands (Giacomello et al., 2020; Mishra and Chan, 2016). Dynamic morphological changes in mitochondria are also observed in plant cells; however, their molecular mechanisms are less understood (Arimura, 2018). A striking morphological transformation of mitochondria is seen during the formation of motile flagellated male gametes termed spermatozoids in bryophytes, which exhibit distinctive morphological characteristics, including two flagella extending from the anterior part of the cell body, a thin and helically elongated nucleus, one plastid located at the posterior side of the nucleus, and two mitochondria, one of which is in the anterior region of the cell body and the other in the posterior region (**Figure 1A**). This mitochondrial distribution differs from that of spermatozoids in other streptophytes and from metazoan spermatozoa (Pitnick et al., 2009; Renzaglia and Garbary, 2001). In spite of morphological investigation of the transformation from immotile spermatids into motile spermatozoids by transmission electron microscopy (TEM), when and how mitochondria are reorganized during spermiogenesis remain mostly unknown.

**Figure 1.**
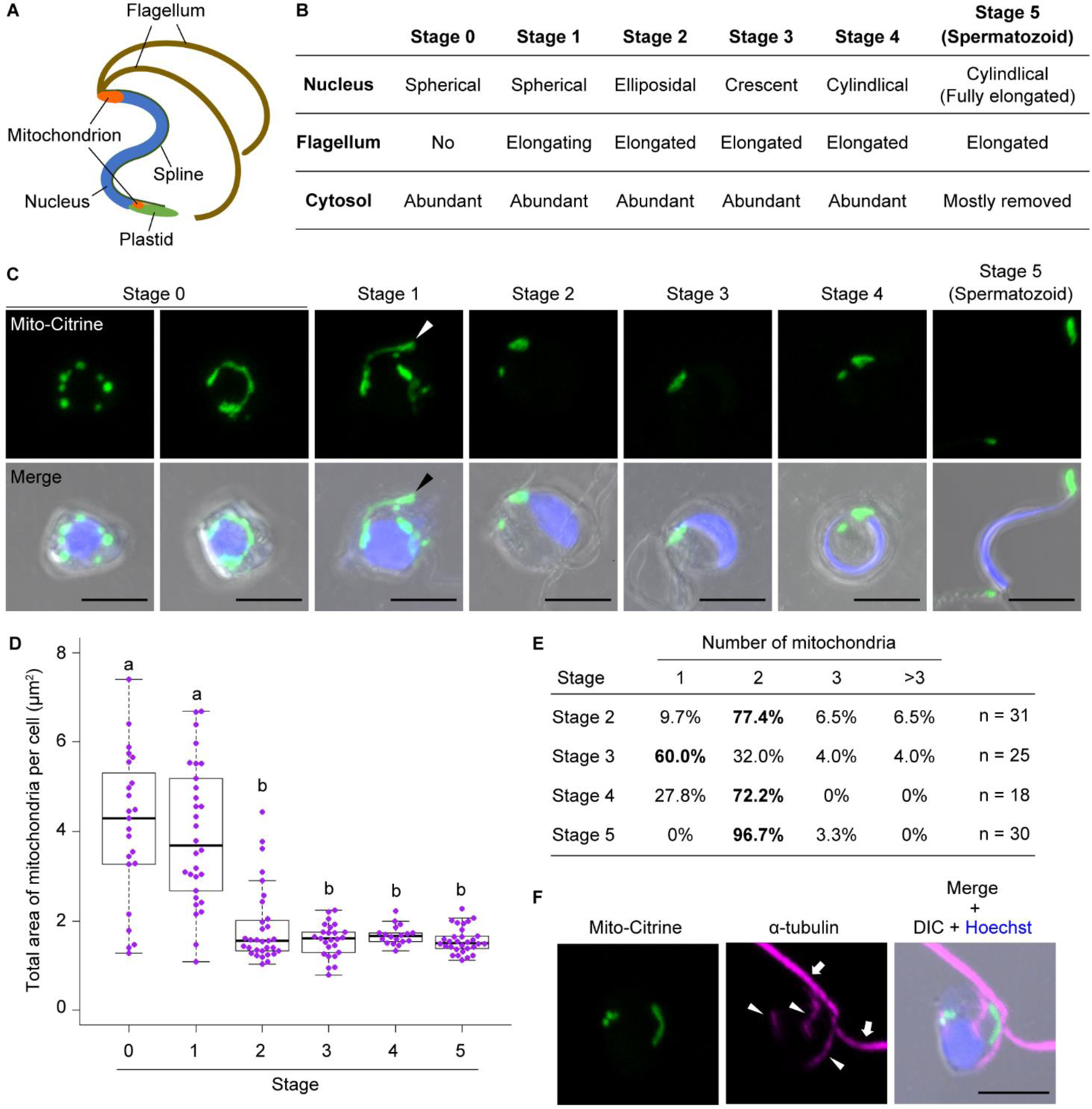
Reorganization of mitochondria during spermiogenesis in *M. polymorpha*. (A) Schematic illustration of a spermatozoid of *M. polymorpha*. (B) Developmental stages (stages 0 to 5) of spermiogenesis in *M. polymorpha* proposed in Minamino et al. (2021). (C) Maximum-intensity projection images of cell wall-digested antheridial cells and a spermatozoid at each developmental stage expressing *_pro_*Mp*EF1α:Mito*-*Citrine* (green). Nuclei were stained with Hoechst 33342 (blue). Arrowheads indicate the elongated mitochondria beneath the base of the flagella. Scale bars = 5 μm. (D) Total area of mitochondria in cells at stages 0 to 5 calculated using maximum-intensity projection images including samples presented in (C). n = 23 (stage 0), 30 (stages 1 and 5), 31 (stage 2), 25 (stage 3), and 18 (stage 4) cells. The boxes and solid lines in the boxes indicate the first quartile and third quartile and the median, respectively. The whiskers indicate 1.5× interquartile ranges. Different letters denote significant differences based on the Steel–Dwass test (*p* < 0.05). (E) The number of mitochondria counted in the same sets of samples analyzed in (D). The bold letter is the most frequent mitochondrial number in each stage. n: the number of cells. (F) Immunostaining of a cell at stage 1 expressing *_pro_*Mp*EF1α:Mito*-*Citrine* (green) using an anti-α-tubulin antibody (magenta). Nuclei were stained by Hoechst 33342 (blue). Arrows and arrowheads indicate flagella and microtubule bundles in the cytoplasm, respectively. Scale bar = 5 μm. DIC: differential interference contrast. See also Figure S1.

Mitochondrial morphology is balanced by mitochondrial fission and fusion. Among the machineries involved in mitochondrial fission, a family of GTPases, dynamin or dynamin-related proteins (DRPs), play essential roles in the constriction and scission of mitochondria (Giacomello et al., 2020). In plant cells, DRP3 has been shown to localize to fission sites on mitochondria and mediate their fission (Arimura and Tsutsumi, 2002; Fujimoto et al., 2009; Nagaoka et al., 2017), whereas the machinery known to act in mitochondrial fusion in nonplant systems has not been detected in plants thus far (Arimura, 2018).

Mitochondrial degradation is another mechanism required for the proper organization of mitochondria in eukaryotic cells, which is mediated partly by autophagy. Among the multiple modes of autophagy reported thus far (Yim and Mizushima, 2020), the involvement of macroautophagy (hereafter referred to as autophagy) in mitochondrial degradation is well documented (Onishi et al., 2021). In autophagy, the double-membrane-bound autophagosome is newly formed from the isolation membrane or the phagophore to engulf and transport cytoplasmic components to the vacuole/lysosome for subsequent degradation (Zhao and Zhang, 2019). Several core autophagy-related (ATG) proteins mediate autophagosome formation and are highly conserved in eukaryotic lineages, including plants (Nakatogawa, 2020; Norizuki et al., 2019; Zhang et al., 2021). Autophagy plays a critical role in bulk degradation of the cytoplasm for nutrient recycling and participates in the selective degradation of various cytoplasmic components (Gatica et al., 2018; Li et al., 2021). Mitochondria are a target of selective degradation by autophagy, which underpins proper mitochondrial functions in various organisms, including plants (Nakamura et al., 2021a; Onishi et al., 2021). Recently, it was reported that spermatids of *atg* mutants of the moss *Physcomitrium patens* comprise a larger number of small mitochondria than wild-type spermatids without a reduction in their total area (Sanchez-Vera et al., 2017). It has been also proposed that a reduction in mitochondrial number is achieved by mitochondrial fusion during spermiogenesis in bryophytes (Renzaglia and Garbary, 2001). These lines of evidence seem to suggest that autophagy is involved in mitochondrial fusion during spermiogenesis, although its precise role and regulation in mitochondrial remodeling remain obscure.

In this study, we first investigated how mitochondria are reorganized during spermiogenesis in the model liverwort *Marchantia polymorpha* (Bowman et al., 2017; Kohchi et al., 2021). We found that mitochondrial morphology changes drastically during spermiogenesis; mitochondria undergo consecutive fission and dissipation at the early stage of spermiogenesis, and only one anterior mitochondrion remains, which then divides to give rise to the posterior mitochondrion. We also found that the unneeded mitochondria are eliminated by autophagy, which precedes autophagic degradation of other cellular components. We further found that some organelles were transported into the vacuole independent of autophagy. These findings indicate that spermiogenesis in *M. polymorpha* occurs through multiple degradation mechanisms, including specifically regulated autophagy, which is essential for the formation of functional spermatozoids.

## RESULTS

### Morphological changes in mitochondria during spermiogenesis in *M. polymorpha*

We started this study by observing alterations in the morphology of mitochondria during spermiogenesis in *M. polymorpha*, which can be divided into 1 + 5 developmental stages based on cellular and nuclear morphology and flagellar formation (**Figure 1B**; Minamino et al., 2021). We visualized mitochondria by expressing Citrine tagged with the mitochondria-targeting signal (Mito-Citrine), whose usability was confirmed by costaining with MitoTracker (**Figure S1A**). At stage 0, two populations of cells were observed; one group possessed many punctate mitochondria, and the other population possessed highly elongated mitochondria surrounding the nucleus (**Figure 1C**). At stage 1, tubular and punctate mitochondria were observed in a cell body, and the tubular (sometimes rod-shaped) mitochondrion was observed at the basement of the flagella (**Figures 1C and 1F**). Since the anterior mitochondrion in plant spermatozoids is associated with basal bodies via the multilayered structure (MLS), which is a microtubule-containing structure unique to plant spermatids and spermatozoids (Kreitner, 1977; Kreitner and Carothers, 1976), this rod-shaped mitochondrion should be the precursor of the anterior mitochondrion. The total area of mitochondria was significantly decreased during stages 1 and 2, and only one mitochondrion remained at stage 3 at the basement of the flagella. At stages 4 and 5, two mitochondria, namely, the anterior and posterior mitochondria, were observed in each cell body (**Figures 1C-1E**). A similar mitochondrial remodeling pattern was also observed when using another mitochondrial marker, MpIDH1-mCitrine (**Figures S1B-S1E**). Thus, the number and size of mitochondria are reduced during early spermiogenesis in *M. polymorpha*, and the posterior mitochondrion is formed by fission of the anterior mitochondrion after the number of mitochondria is reduced to one at stage 3.

### The posterior mitochondrion is formed by MpDRP3-dependent fission of the anterior mitochondrion

We attempted to verify whether mitochondrial fission occurs during mitochondrial remodeling during spermiogenesis in *M. polymorpha*. A dynamin-related protein encoded by MpDRP3 plays a critical role in mitochondrial fission in *M. polymorpha* (Nagaoka et al., 2017). We expressed 3×HA (human influenza hemagglutinin)-tagged MpDRP3l, the product of the longer transcriptional variant of Mp*DRP3* transcripts (Nagaoka et al., 2017), under the regulation of its own promoter. In this plant, we detected the subcellular localization of 3×HA-MpDRP3l by immunostaining with an anti-HA antibody in spermatids undergoing spermiogenesis. At stages 0 and 1, 3×HA-MpDRP3l was detected as foci on mitochondria in spermatids, suggesting that mitochondrial fission occurs frequently at these stages (**Figure 2A**). However, at stage 3, the signal from 3×HA-MpDRP3l was hardly detected in spermatids, probably reflecting that mitochondrial fission scarcely occurs at this stage. This result is consistent with the result that punctate mitochondria were frequently observed during stages 0 to 1, but only one mitochondrion was observed at stage 3 (**Figure 1**).

**Figure 2.**
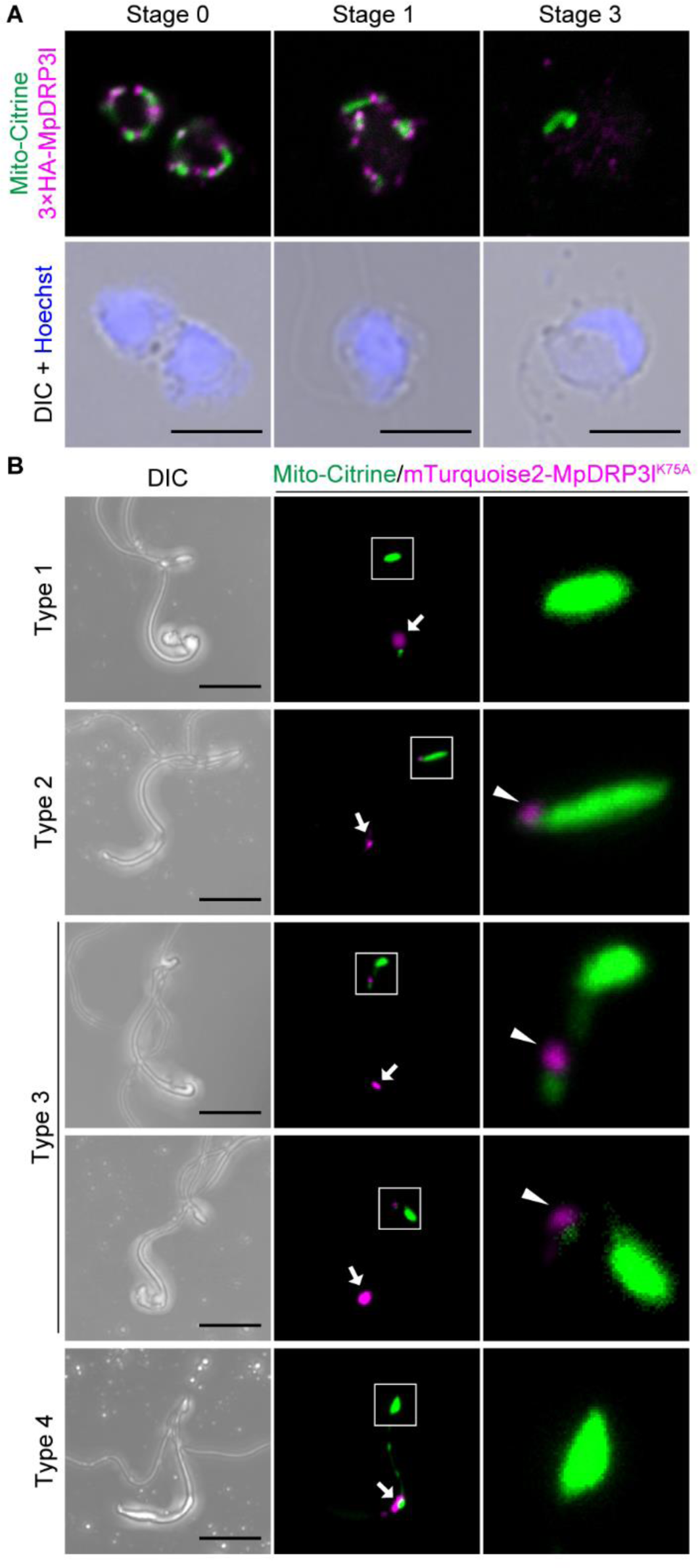
MpDRP3-mediated mitochondrial fission in mitochondrial reorganization during spermiogenesis. (A) Confocal images of antheridial cells at stages 0, 1, and 3 expressing *_pro_*Mp*EF1α:Mito-Citrine* (green) and *_pro_*Mp*DRP3:3×HA*-Mp*DRP3l* (magenta). Nuclei were stained with Hoechst 33342 (blue). Scale bars = 5 μm. (B) Maximum-intensity projection images of spermatozoids with the indicated types of mitochondria expressing *_pro_*Mp*EF1α:Mito*-*Citrine* (green) and *_pro_*Mp*MS1:mTurquoise2-*Mp*DRP3l^K75A^* (magenta). Nuclei were stained with Hoechst 33342 (blue). Arrows indicate MpDRP3l^K75A^ in the posterior region of spermatozoids and arrowheads show MpDRP3l^K75A^ associated with anterior mitochondria. Right panels are magnified images of the boxed regions. Scale bars = 5 μm. See also Figure S2.

We then examined whether mitochondrial fission is a prerequisite for mitochondrial reduction during spermiogenesis. The Mp*drp3* knockout mutant exhibits a severe defect in thallus growth (Nagaoka et al., 2017); therefore, it was not practical to see the effect of this mutation after sexual organ formation. Here, we took advantage of a dominant negative form of MpDRP3l to inhibit the function of MpDRP3 specifically during spermatogenesis. Dynamin family members, whose conserved lysine residue in the GTPase domain is replaced with alanine, exhibit no GTPase activity and exert dominant-negative effects (**Figure S2A**; Arimura et al., 2002; Naylor et al., 2006). We expressed the mutant version of MpDRP3l (MpDRP3l^K75A^) tagged with mTurquoise2 under the regulation of the Mp*MS1* promoter, which is predominantly active in antheridia (Higo et al., 2016), and examined its effect on mitochondrial fission during spermiogenesis. At the early stage of spermiogenesis (stage 1), we frequently observed only one elongated tubular mitochondrion in the cell body of spermatids, in contrast to wild-type spermatids that harbored punctate mitochondria in addition to elongated mitochondria (**Figures S2B and S2C**). We also examined the effect of the expression of wild-type MpDRP3l tagged with mTurquoise2 and detected no alteration in the ratio of cells with one elongated mitochondrion (**Figures S2B and S2C**). These results indicated that the mutant version of MpDRP3l inhibited mitochondrial fission in a dominant-negative manner, as reported previously. However, spermatozoids expressing the mutant MpDRP3l were not distinguishable from wild-type spermatozoids by shape when observed by light microscopy (**Figure S2D**). We then observed mitochondria in spermatozoids. Intriguingly, the total area of mitochondria in each spermatozoid expressing MpDRP3l^K75A^ was not significantly increased compared with that in wild-type spermatids, suggesting that defective mitochondrial fission did not markedly affect the elimination of mitochondria (**Figure S2F**).

However, the morphology of mitochondria in spermatozoids was significantly altered by the expression of mutant MpDRP3l (**Figures 2B, S2D and S2E**). We classified the mitochondrial morphology in spermatozoids into four types: spermatozoids with one anterior and one posterior mitochondrion, such as wild-type spermatozoids (type 1); spermatozoids with only one anterior mitochondrion (type 2); spermatozoids with two or three closely associated (sometimes seemingly connected) mitochondria in their anterior region (type 3); and spermatozoids with additional fluorescent spots besides the anterior and posterior mitochondria (type 4) (**Figures 2B, S2D and S2E**). Type 2 mitochondria were frequently associated with mTurquoise2-MpDRP3l^K75A^ at the posterior tip (85.5% of 69 spermatozoids), and fluorescence from this protein was observed to be associated with 87.5% of anterior mitochondria in type 3 spermatozoids (n =16 spermatozoids), 57.1% of which showed MpDRP3l^K75A^ accumulation between two neighboring anterior mitochondria (**Figure 2B**). A strong signal from mTurquoise2 was also observed in the posterior region without association with any mitochondria for all types of spermatozoids, which probably represents the aggregated protein in the remnant of the cytosol (arrows in **Figure 2B**). Abnormal mitochondria of types 2 to 4 were rarely observed in spermatids expressing wild-type MpDRP3l (**Figure S2E**). These results indicated that the posterior mitochondrion is formed by MpDRP3-dependent fission of the anterior mitochondrion.

### Autophagy plays essential roles in normal spermiogenesis

In addition to mitochondria, a major part of the cytoplasm, including endomembrane organelles, is also eliminated from the cell body during spermiogenesis (Minamino et al., 2017; Minamino et al., 2021). This drastic organelle remodeling during spermiogenesis could be completed through activated degradation pathways such as autophagy, therefore, we investigated whether mutants defective in autophagy exhibit any defects in spermiogenesis. In contrast to wild-type spermatozoids with elongated helical cell bodies containing thin cylindrical nuclei with trace amounts of cytoplasm, spermatozoids of Mp*atg5-1^ge^*, which was generated using the CRISPR/Cas9 system (Norizuki et al., 2019), harbored abundant cytoplasm and abnormally shaped nuclei that were significantly more circular than wild-type spermatozoids (**Figures 3A-3C**).

**Figure 3.**
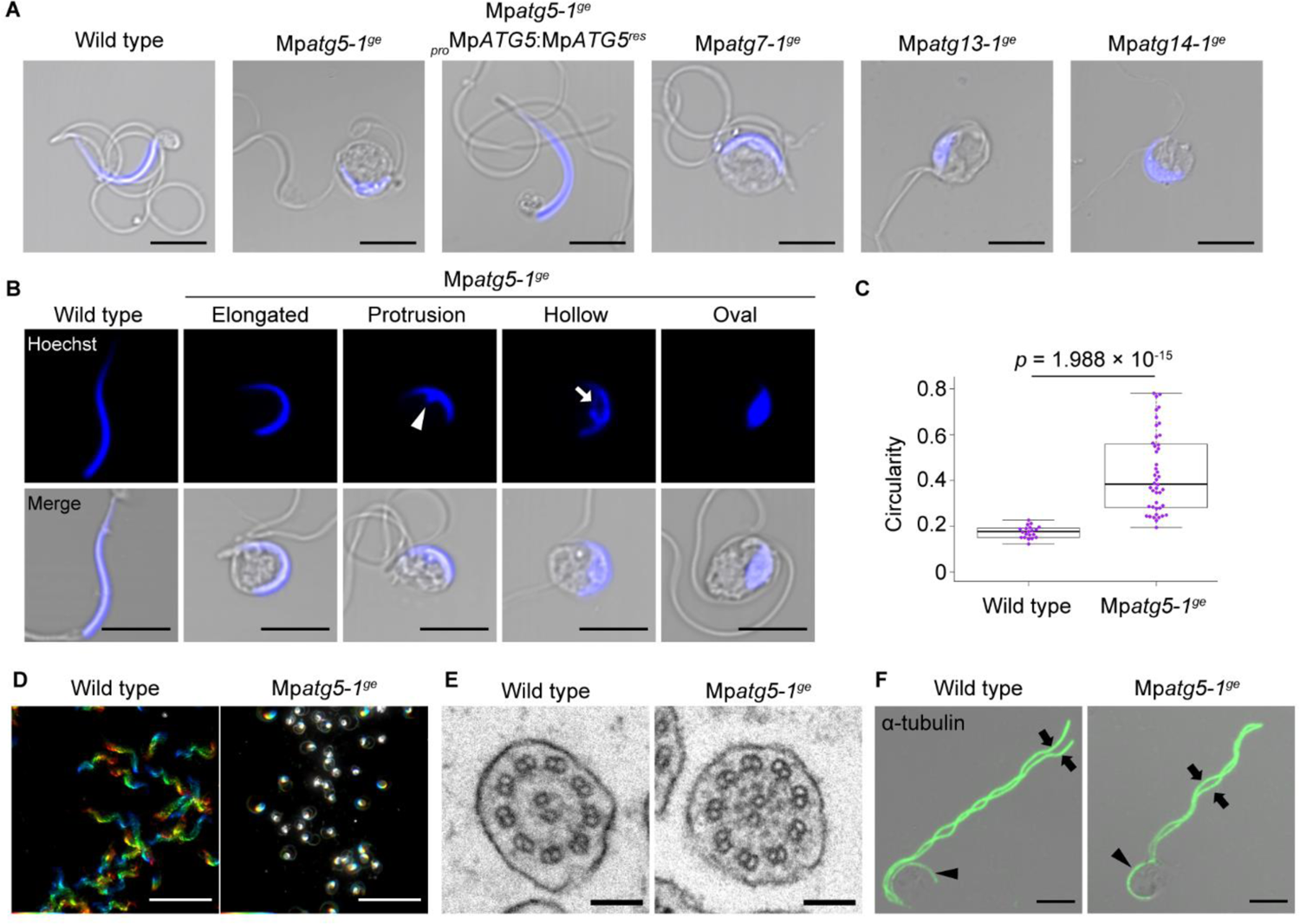
Abnormal spermatozoids formed in mutants of core *ATG* genes. (A) Confocal images of wild-type and Mp*atg* mutant spermatozoids. Nuclei were stained with Hoechst 33342 (blue). Scale bars = 5 μm. (B) Confocal images of wild-type and Mp*atg5-1^ge^* spermatozoids. Nuclei were stained with Hoechst 33342 (blue). The arrowhead and arrow indicate the protrusion and hollow of the nucleus, respectively. Scale bars = 5 μm. (C) Circularity of the nucleus calculated using samples whose nuclei were stained with Hoechst 33342, as shown in (B). n = 20 (wild-type) and 44 (Mp*atg5-1^ge^*) spermatozoids. The *p* value obtained by the Wilcoxon rank sum test is shown. (D) Trajectories of movement of wild-type and Mp*atg5-1^ge^* spermatozoids. Thirty-three images taken in one second were differently pseudocolored and overlayed. Scale bars = 50 μm. (E) TEM images of axonemes in flagella in wild-type and Mp*atg5-1^ge^* spermatids. Scale bars = 100 nm. (F) Immunostaining of wild-type and Mp*atg5-1^ge^* spermatids with the anti-α-tubulin antibody (green). Arrows and arrowheads indicate flagella and the spline, respectively. Scale bars = 5 μm. See also Figures S3 and S4.

The mutant spermatozoids also exhibited defects in motility and fertility (**Figures 3D and S4**). However, despite impaired motility, we did not detect any marked structural differences in the flagellar axoneme or spline between wild-type and Mp*atg5-1^ge^* spermatids (**Figures 3E and 3F**). All of the abnormalities were restored by introducing CRISPR/Cas9-resistant Mp*ATG*5 (Mp*ATG5^res^*), confirming that these defects resulted from the loss of function of Mp*ATG5* (**Figure 3A**). Similar phenotypes to those in Mp*atg5-1^ge^* were also observed in the mutant of Mp*ATG7* (Mp*atg7^ge^*; Norizuki et al., 2019), which acts in lipidation of MpATG8 together with Mp*ATG5*, and mutants of Mp*ATG13* and Mp*ATG14* (Mp*atg13^ge^* and Mp*atg14^ge^*) defective in distinct steps in autophagosomal formation from Mp*ATG5*, further demonstrating that the abnormalities we detected were attributed to defective autophagy (**Figures 3A, S3, and S4**).

**Figure 4.**
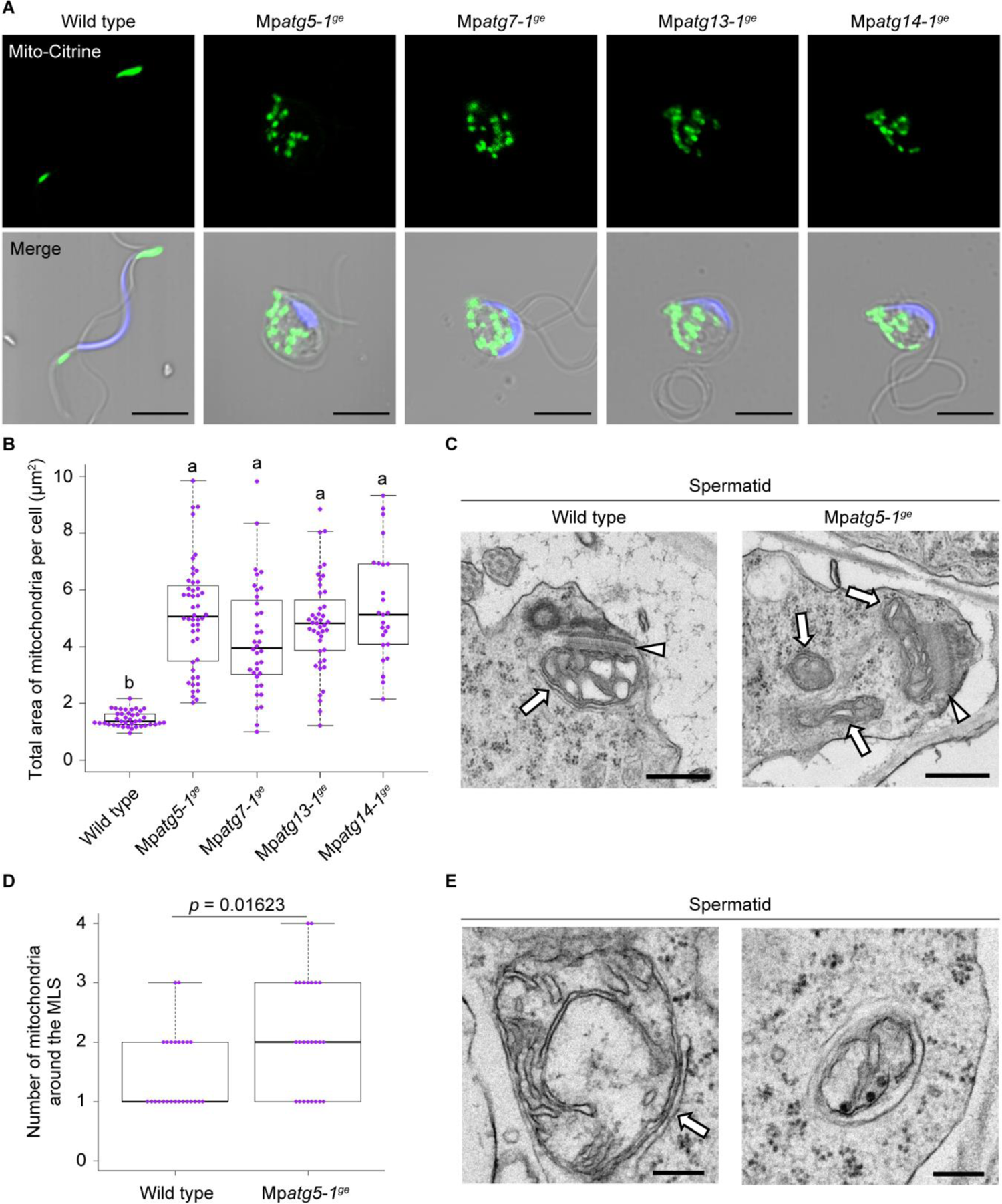
Impaired reorganization of mitochondria during spermiogenesis in autophagy-defective mutants. (A) Confocal images of wild-type and Mp*atg* mutant spermatozoids expressing *_pro_*Mp*EF1α:Mito*-*Citrine* (green). Nuclei were stained with Hoechst 33342 (blue). Scale bars = 5 μm. (B) The total area of mitochondria in wild-type and Mp*atg* mutant spermatozoids. The mitochondrial area was calculated in maximum-intensity projection images of samples including spermatozoids presented in (A). n = 40 (wild-type), 48 (Mp*atg5-1^ge^*), 36 (Mp*atg7-1^ge^*), 43 (Mp*atg13-1^ge^*), and 25 (Mp*atg14-1^ge^*) spermatozoids. The boxes and solid lines in the boxes indicate the first quartile and third quartile and the median, respectively. The whiskers indicate 1.5× interquartile ranges. Different letters denote significant differences based on the Steel–Dwass test (*p* < 0.05). (C) TEM images of wild-type and Mp*atg5-1^ge^* spermatids. Arrows and arrowheads indicate mitochondria and the MLS, respectively. Scale bars = 500 nm. (D) The number of mitochondria around the MLS counted in TEM images taken at ×21,100, including those presented in (C). n = 25 images for each genotype. Statistical differences were analyzed by the Wilcoxon rank sum test. (E) TEM images of mitochondria in wild-type spermatids. In the left panel, a mitochondrion was associated with the isolation membrane-like structure (arrow). In the right panel, a mitochondrion was engulfed seemingly selectively by the autophagosome. Scale bars = 200 nm. See also Figure S5.

### Autophagy is required for mitochondrial reorganization during spermiogenesis

Autophagy is involved in the degradation of damaged and/or excess mitochondria in various organisms (Nakamura et al., 2021a; Onishi et al., 2021). We then investigated mitochondria in Mp*atg* spermatids and spermatozoids to investigate whether mitochondria in spermatids are degraded by autophagy. While wild-type spermatozoids possessed two mitochondria, Mp*atg* spermatozoids possessed many punctate and/or rod-shaped mitochondria (**Figures 4A, 4B, S5A, and S5B**). We then performed TEM observation around the anterior mitochondrion, which is discernable by association with the MLS, which confirmed that a larger number of mitochondria other than the anterior mitochondria remained in the Mp*atg5-1^ge^* spermatids than in wild-type spermatids (**Figures 4C and 4D**).

These results suggested that mitochondrial degradation during spermiogenesis is impaired in autophagy-deficient mutants. While Mp*atg* spermatozoids accumulated fragmented mitochondria, which occasionally formed clusters, elongated mitochondria were observed in spermatids of Mp*atg5-1^ge^* at stages 0 and 1, as in the wild type, suggesting that mitochondrial fission is not impaired by the Mp*atg* mutations (**Figure S5C**). During TEM observation of wild-type spermatids undergoing spermiogenesis, we also observed mitochondria associated with the isolation membrane-like structure, which should be the precursor of the autophagosome, and a mitochondrion seemingly selectively engulfed by the autophagosome (**Figure 4E**). These structures implied that mitochondria are selectively degraded by autophagy in spermatids rather than by bulk autophagy.

### Autophagy is also involved in the reorganization of other organelles but is distinct from the autophagy of mitochondria

Because the Mp*atg* mutants were defective in cytoplasm removal during spermiogenesis, autophagy could also be involved in the elimination of organelles other than mitochondria. To verify this possibility, we then observed the endoplasmic reticulum (ER) and the Golgi apparatus in Mp*atg5-1^ge^* spermatozoids. In the mutant spermatozoids, the ER marker mCitrine-MpSEC22 (Kanazawa et al., 2016) was retained in the cytoplasm, whereas this marker was not detected in wild-type spermatozoids (**Figure 5A**). We then observed two Golgi markers, mCitrine-MpGOS11 and ST-Venus (Kanazawa et al., 2016), in spermatozoids. Intriguingly, these Golgi markers were distinctly affected by the Mp*atg5-1^ge^* mutation. In wild-type spermatozoids, neither of these markers was detected in spermatozoids, as reported previously (Minamino et al., 2021). In Mp*atg5-1^ge^* spermatozoids, mCitrine-MpGOS11 was not detected in the cell body (**Figure 5A**), indicating that the removal of MpGOS11 does not require MpATG5. Conversely, ST-Venus remained in the cytoplasm of Mp*atg5-1^ge^* spermatozoids (**Figure 5A**). These results indicated that the degradation of Golgi proteins distinctly involves autophagy, although the Golgi apparatus is eventually removed completely from wild-type spermatozoids.

**Figure 5.**
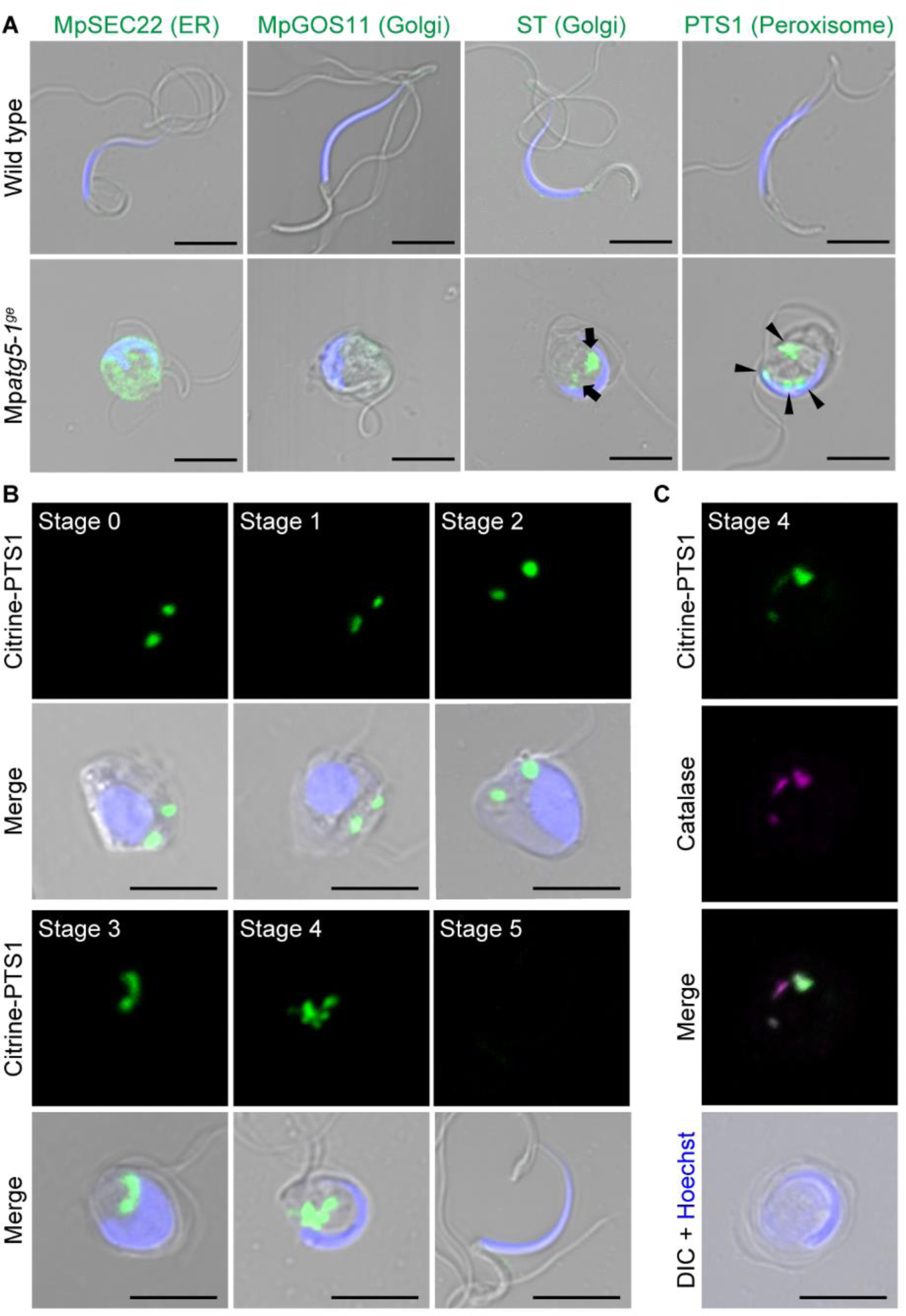
Impaired elimination of the ER, Golgi apparatus, and peroxisomes during spermiogenesis in autophagy-defective mutants. (A) Confocal images of wild-type and Mp*atg5-1^ge^* spermatozoids expressing the ER marker *_pro_*Mp*DUO1:mCitrine*-Mp*SEC22*, the Golgi marker *_pro_*Mp*DUO1:mCitrine*-Mp*GOS11* or *_pro_*Mp*DUO1:ST*-*Venus*, and the peroxisome marker *_pro_*Mp*EF1α:Citrine*-*PTS1,* which are shown in green. Nuclei were stained with Hoechst 33342 (blue). Arrows and arrowheads indicate the fluorescence from ST-Venus and Citrine-PTS1, respectively. Scale bars = 5 μm. (B) Confocal images of cell wall-digested antheridial cells and spermatozoids expressing the peroxisome marker *_pro_*Mp*EF1α:Citrine*-*PTS1* (green). Nuclei were stained with Hoechst 33342 (blue). Scale bars = 5 μm. (C) Wild-type spermatids at stage 4 expressing *_pro_*Mp*EF1α:Citrine*-*PTS1* (green). Catalase was detected by immunostaining (magenta), and nuclei were stained with Hoechst 33342 (blue). Scale bar = 5 μm.

We then examined whether peroxisomes are also removed during spermiogenesis in *M. polymorpha* using a peroxisome marker, Citrine-PTS1 (Mano et al., 2018). In wild-type spermatozoids, the signal from Citrine-PTS1 was not detected. However, in Mp*atg5-1^ge^* spermatozoids, Citrine-PTS1 was observed as multiple puncta, suggesting that autophagy is required for the removal of peroxisomes during spermiogenesis (**Figure 5A**). We then investigated when Citrine-PTS1 disappeared during spermiogenesis in the wild type and found that it remained detectable as puncta through stage 0 to stage 4 (**Figure 5B**). This signal from Citrine-PTS1 represented peroxisomes but not cytosolic aggregation, because catalase was detected in the puncta by immunostaining using an anti-catalase antibody (Mano et al., 2018) (**Figure 5C**). Thus, peroxisome degradation in spermatids requires autophagy and occurs between stages 4 and 5. Notably, peroxisome degradation occurs at a different stage from mitochondrial degradation, which is mostly completed by stage 2 (**Figures 1C-1E**). This finding indicates that mitochondria and peroxisomes are removed by autophagy at different stages, suggesting a highly regulated and elaborate mechanism of autophagy for degrading organelles during spermiogenesis in *M. polymorpha*.

### Distinct requirement of MpATG5 in organelle degradation

The defective removal of multiple organelles during spermiogenesis in Mp*atg5-1^ge^* implied that those organelles could be transported to and degraded in the vacuole via autophagy. Meanwhile, we demonstrated that two Golgi marker proteins are distinctly affected by the Mp*atg5-1^ge^* mutation, suggesting that degradation mechanisms other than MpATG5-dependent autophagy also mediate spermiogenesis in *M. polymorpha*. For more information on the vacuolar degradation of organelles, we then examined the accumulation of organelle markers in vacuoles by imaging the vacuolar membrane and organelle markers in spermatids. For this experiment, the Mp*atg5-1^ge^* mutant without the T-DNA comprising elements for genome editing (Mp*atg5-1^ge/cf^*) was generated (**Figure S6**), and the vacuolar membrane marker, XFP-tagged MpVAMP71, and fluorescently tagged organelle markers were introduced.

We first confirmed that the autophagosome marker mTurquoise2-MpATG8a, which is transported into the spherical vacuole in wild-type spermatids undergoing spermiogenesis, was not detected in the spherical vacuole in Mp*atg5-1^ge/cf^* (**Figure 6A**), consistent with the results in thallus cells (Norizuki et al., 2019). We next observed vacuolar transport of the plasma membrane (PM) marker mCitrine-MpSYP12A (Kanazawa et al., 2016), which should be transported via the endocytic pathway but not by autophagy, and found that a signal from mCitrine was detected in the spherical vacuole in both wild-type and Mp*atg5-1^ge/cf^* spermatids (**Figure 6B**). This result also confirmed that the mutation in Mp*ATG5* did not markedly affect the endocytic pathway for the vacuole.

**Figure 6.**
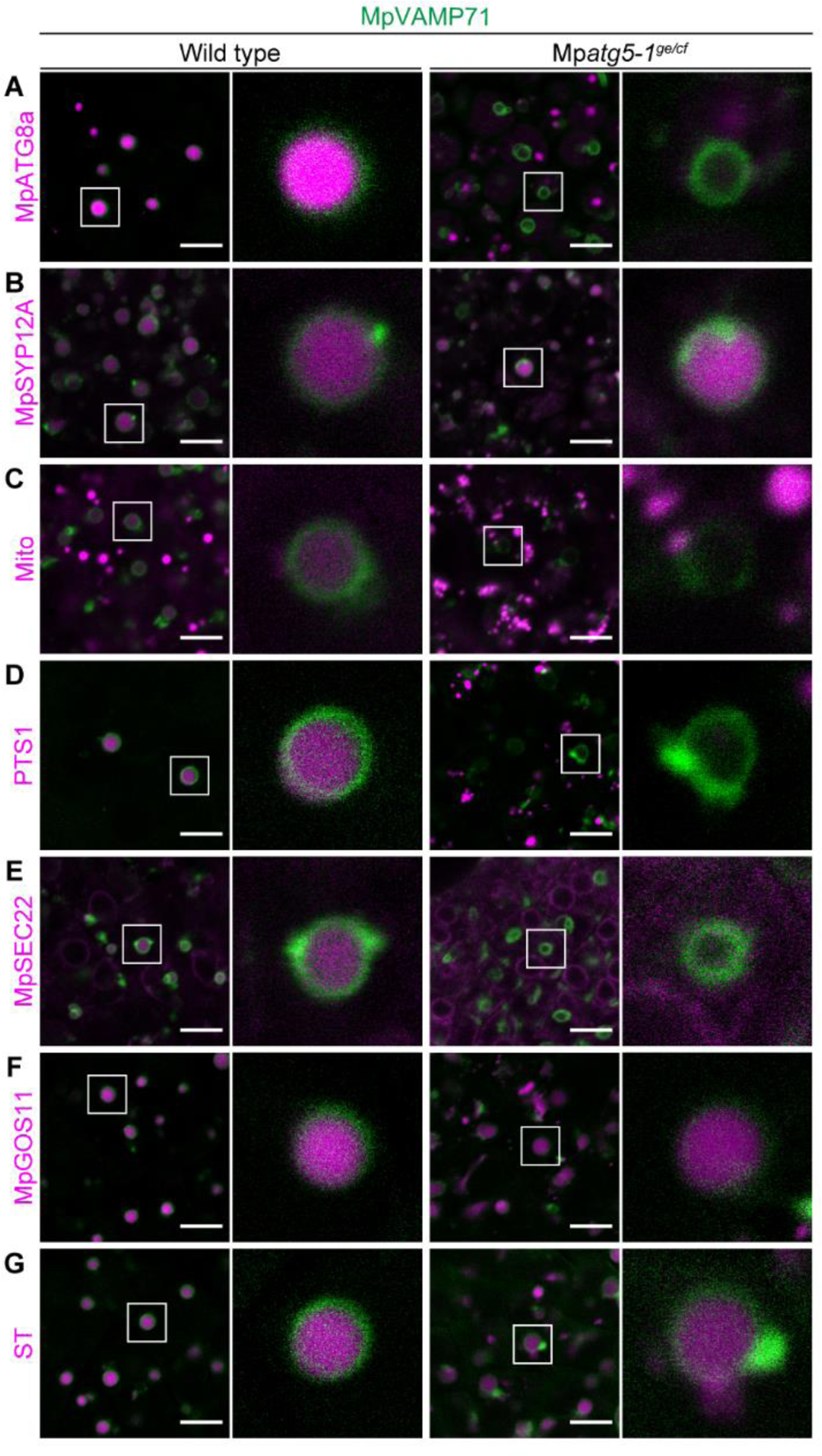
Some organelle proteins are transported into the vacuole independently of MpATG5 during spermiogenesis Confocal images of vacuolar membrane markers (fluorescently tagged MpVAMP71, green) and other organelle markers (magenta) in hand-sectioned wild-type and Mp*atg5-1^ge/cf^* antheridial cells. The autophagosome marker *_pro_*Mp*ATG8a:mTurquoise2*-Mp*ATG8a* (A), the PM marker *_pro_*Mp*SYP12A:mCitrine*-Mp*SYP12A* (B), the mitochondrial marker *_pro_*Mp*SYP2:Mito-Citrine* (C), the peroxisome marker *_pro_*Mp*EF1α:mTurquoise2*-*PTS1* (D), the ER marker *_pro_*Mp*DUO1:mCitrine*-Mp*SEC22* (E), and the Golgi marker *_pro_*Mp*DUO1:mTurquoise2*-Mp*GOS11* (F) or *_pro_*Mp*DUO1:ST*-*mTurquoise2* (G) were expressed with *_pro_*Mp*VAMP71:mCitrine*-Mp*VAMP71* (A, D, F, and G) or *_pro_*Mp*VAMP71:mGFP*-Mp*VAMP71* (B, C, and E). Distinct from other experiments, Mito-Citrine is expressed under the regulation of the Mp*SYP2* promoter at a moderate expression level. Scale bars = 5 μm. See also Figure S6.

Next, we investigated whether the transport of other organelle markers to the vacuole requires MpATG5-dependent autophagy. Consistent with the phenotype in the Mp*atg5-1^ge^* spermatozoid, markers for mitochondria, peroxisomes, and the ER were not detected in the spherical vacuole in Mp*atg5-1^ge/cf^*, although they accumulated in the vacuole in wild-type spermatids (**Figures 6C-6E**). This result confirmed that both the mitochondria and peroxisomes are transported to the vacuole only by autophagy, although these organelles are degraded in distinct developmental stages during spermiogenesis. Intriguingly, the Golgi markers mTurquoise2-MpGOS11 and ST-mTurquoise2 were transported into the vacuole even in Mp*atg5-1^ge/cf^* spermatids (**Figures 6F and 6G**), although the removal of ST-Venus from the cytoplasm required MpATG5 activity (**Figure 5A**). This result suggested that a part of the Golgi apparatus, or at least some part of the Golgi-resident proteins, could be transported into the vacuole in an MpATG5-independent manner, although complete clearance of the Golgi apparatus requires autophagy. In summary, autophagy distinctly contributes to the removal of each organelle, is tightly regulated during spermiogenesis, and cooperates with nonautophagic degradation pathways to accomplish spermiogenesis in *M. polymorpha* (**Figure 7**).

**Figure 7.**
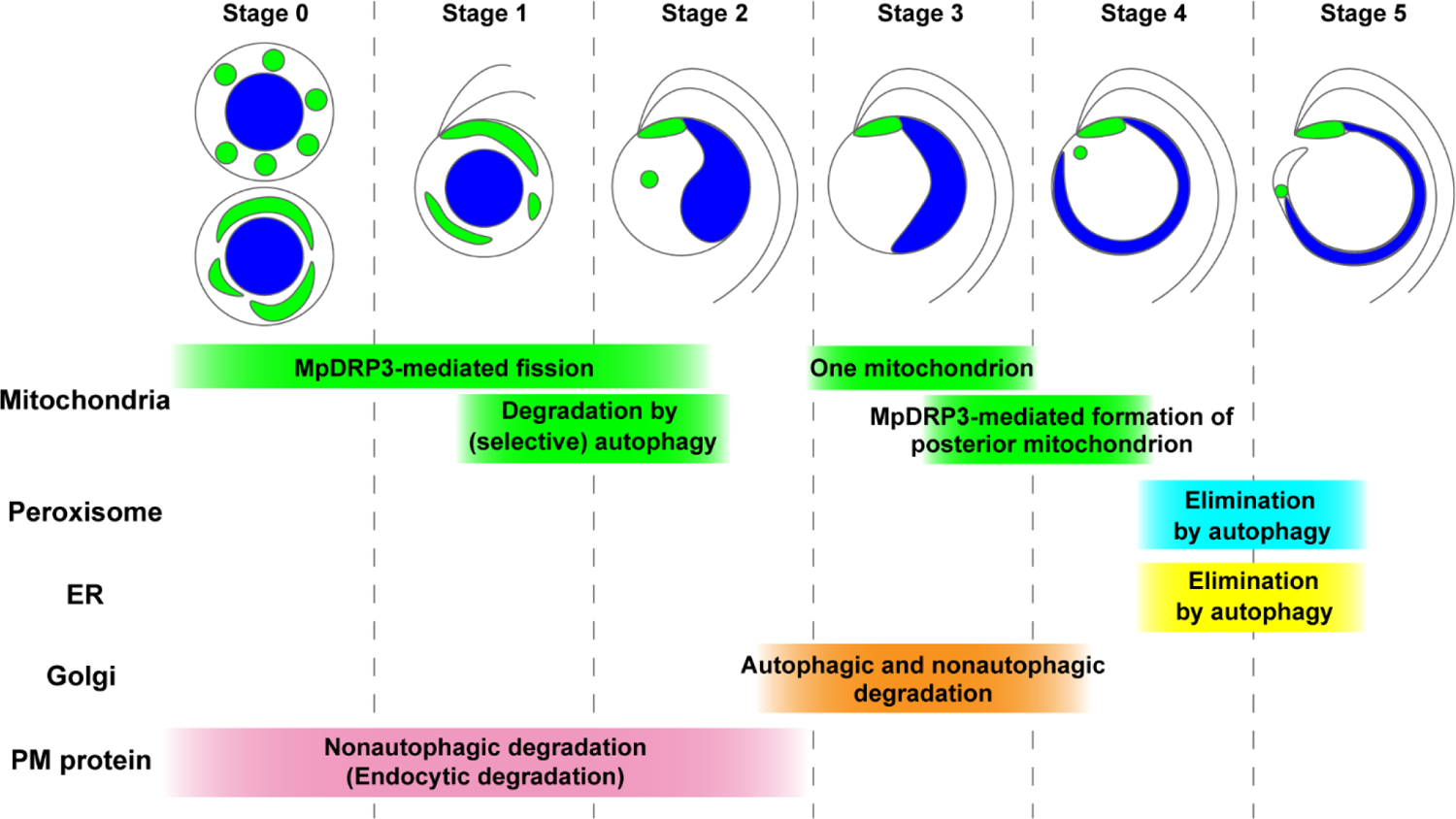
Schematic diagram of organelle reorganization during spermiogenesis in *M. polymorpha* Mitochondria undergo MpDRP3-mediated fission and degradation by (selective) autophagy during stages 0 to 2. One mitochondrion remains at the base of the flagella in stage 3, from which the posterior mitochondrion is formed through MpDRP3-dependent fission, resulting in two mitochondria in a spermatid. While the removal of unneeded mitochondria is mostly accomplished by stage 2, autophagic degradation of peroxisomes and the ER occurs after stage 4. Elimination of the Golgi apparatus involves autophagic and nonautophagic degradation, which occurs around stage 3. The PM is drastically remodeled during stages 0 to 2; MpSYP12A is removed from the PM by endocytosis and degraded in the vacuole independently of autophagy. Upper illustrations show morphological changes in the mitochondria (green), the nucleus (blue), and the flagella. We referred to Minamino et al. (2021) for the stages of removal of the PM protein (MpSYP12A), the ER, and the Golgi apparatus.

## DISCUSSION

### Drastic reorganization of mitochondria during spermiogenesis in *M. polymorpha*

Mitochondrial morphology is reported to be closely related to mitochondrial functions in various organisms. For instance, elongated mitochondria are bioenergetically efficient and therefore observed in conditions that require increased ATP production (Giacomello et al., 2020; Mishra and Chan, 2016). We found that highly elongated mitochondria, which were not observed in thallus cells grown under our normal conditions, were observed in cells at stages 0 and 1 (**Figures 1 and S1**). Given that *de novo* synthesis of flagella occurred between stages 0 and 2 (Minamino et al., 2021), the elongated mitochondria at these stages might reflect high energy demand at these stages.

The localization of MpDRP3l on mitochondria during stages 0 and 1 and discrete punctate or rod-shaped morphology of mitochondria during stage 1 (**Figures 1C and 2A**), together with the elongated morphology induced by expression of MpDRP3l^K75A^ (**Figures S2B and S2C**), strongly suggested that mitochondria underwent MpDRP3-dependent fission during these stages. Intriguingly, small punctate mitochondria disappeared before stage 3, and only one rod-shaped mitochondrion was detected at the base of the flagella at this stage (**Figures 1C and 1F**). Anterior mitochondrion is located at the base of the flagella, which is associated with the MLS, and basal bodies are attached to the uppermost layer of the MLS (spline) (Kreitner, 1977; Kreitner and Carothers, 1976). The similar localization of the rod-shaped mitochondrion at the base of the flagella at stage 3 would indicate that this mitochondrion should be the precursor of the anterior mitochondrion. We demonstrated that the number of mitochondria was reduced to one at stage 3 (**Figure 1**), although the monomitochondrion stage has not been observed during liverwort spermiogenesis. Although we did not detect the accumulation of 3×HA-MpDRP3l at the fission site of the precursor mitochondria during stages 3 to 4, probably due to its transient nature, MpDRP3 was suggested to be involved in the formation of the posterior mitochondrion by two results; 1) the majority of spermatozoids expressing MpDRP3l^K75A^ failed to form the posterior mitochondrion, and 2) MpDRP3l^K75A^ was observed to accumulate on the anterior mitochondrion in type 2 and type 3 spermatozoids that lacked the posterior mitochondrion (**Figures 2B, S2D, and S2E**). MpDRP3l^K75A^ expression also increased the proportion of type 4 spermatozoids, which appeared to possess more than two fluorescent domains, as shown by Mito-Citrine. Additional fluorescent spots were observed between the anterior and posterior mitochondria, and MpDRP3l^K75A^ caused elongation of mitochondria when expressed in spermatids undergoing early spermiogenesis. Thus, these fluorescent foci of Mito-Citrine in type 4 spermatozoids might represent subregions in thin tubular mitochondria continuous with the anterior and posterior mitochondria, whose matrix space other than the fluorescent regions was too narrow to include Mito-Citrine at a detectable level. Otherwise, MpDRP3l^K75A^ might promote mitochondrial fission through an unknown mechanism.

### Mitochondria are degraded by autophagy during spermiogenesis

In this study, we showed that mitochondria are degraded by autophagy, except for one specific mitochondrion, which is the precursor of the anterior and posterior mitochondria, although we do not rule out the possibility that mitochondrial fusion also contributes to a reduction in the mitochondrial number in *M. polymorpha* as proposed previously (Renzaglia and Garbary, 2001). Although mitochondrial fission was inhibited by expression of MpDRP3l^K75A^ at early spermiogenesis, the total size of mitochondria in spermatozoids was not significantly affected, suggesting that MpDRP3-mediated mitochondrial fission during early spermiogenesis is not essential for the removal of mitochondria by autophagy (**Figure S2**). In non-plant systems, the autophagic degradation of mitochondria independent of dynamin members has also been reported (Burman et al., 2017; Yamashita et al., 2016). It would be an interesting future project to study the physiological significance of mitochondrial fission during early spermiogenesis in *M. polymorpha*.

Our results demonstrated that mitochondria are removed at a distinct time from other organelles during spermiogenesis; i.e., the majority of mitochondria were degraded in stage 1, whereas peroxisomes remained at stage 4 (**Figure 7**). The ER, Golgi apparatus, vacuole, and cytosol are also removed from the spermatids later than the mitochondria (Minamino et al., 2021). This result could be explained by the idea that mitochondria are selectively degraded during early spermiogenesis, distinct from other organelles. Our observation that mitochondria appeared to be selectively engulfed by autophagosomes (**Figure 4E**) supported this notion. It is also interesting to ask how one mitochondrion is selected for protection from degradation. In *Arabidopsis thaliana* and metazoa, damaged mitochondria are degraded through selective autophagy, termed mitophagy (Ma et al., 2021; Nakamura et al., 2021b; Narendra et al., 2008). However, we did not find abnormalities in the morphology of mitochondria remaining in Mp*atg5-1^ge^* spermatids by TEM observation (**Figure 4C**), suggesting that damage to mitochondrial function would not be the cue for autophagic degradation in spermatids undergoing spermiogenesis. Although the molecular mechanism of mitophagy has been intensively studied in non-plant systems, it remains largely unknown in plants (Nakamura et al., 2021a). Further studies are needed to unravel the molecular mechanism of mitophagy in plants, and the spermiogenesis of *M. polymorpha* would be a useful system for it.

### Multimode degradation systems mediate spermiogenesis in *M. polymorpha*

We demonstrated that autophagy is also required for the reorganization of organelles other than mitochondria. Autophagy was responsible for the clearance of the ER and peroxisomes (**Figures 5A and 6**), although it remains unclear whether these organelles are removed by selective or nonselective autophagy. Of the markers for the Golgi apparatus, ST-Venus but not mCitrine-MpGOS11 remained in the Mp*atg5-1^ge^* spermatozoid, suggesting that a part of the Golgi apparatus or Golgi proteins are degraded by MpATG5-dependent autophagy during spermiogenesis (**Figure 5A**). Intriguingly, however, ST-mTurquoise2 was also detected in the spherical vacuole in Mp*atg5-1^ge/cf^* spermatids, which indicated that a subpopulation of ST-mTurquoise2 was also transported into the vacuole independently of MpATG5 (**Figure 6G**). The endosomal sorting complex required for transport (ESCRT)-dependent transport of membrane proteins into the vacuole should be highly active in *M. polymorpha* spermatids undergoing spermiogenesis, given that PM proteins such as MpSYP12A are rapidly internalized and degraded in the vacuole during spermiogenesis (**Figure 6B**; Minamino et al., 2017), which should not require autophagy. It has also been reported that microautophagy-like transport, which is accomplished by the vacuolar membrane engulfing substrates independently of ATG5, occurs in *A. thaliana* (Chanoca et al., 2015). Another mechanism for the degradation of cytoplasmic components, the Golgi membrane-associated degradation pathway, has also been reported in non-plant systems (Yamaguchi et al., 2016). One or more of these pathways might contribute to transporting ST-Venus/ST-mTurquoise2 to the vacuole, which should be verified in future studies. Another interesting question is how the vacuole is eliminated after autophagic and nonautophagic degradation of other organelles. The large spherical vacuole remains in the cell body by stage 4, which should be eliminated from the spermatid for completion of spermiogenesis. Investigation of the mechanism of the vacuole removal should be also conducted in future studies.

Autophagy is involved in male gametogenesis in mammals and bryophytes (Norizuki et al., 2020), but their roles seem to diverge. Mice defective in autophagy exhibit defects in the reorganization of microtubules and organelles, and similar effects have also been reported in the moss *P. patens* (Huang et al., 2021; Sanchez-Vera et al., 2017; Shang et al., 2016; Wang et al., 2014). However, in *M. polymorpha*, autophagy does not seem to be involved in microtubule organization; we did not detect alterations in axoneme and spline structures, in contrast to the observations in *P. patens* (**Figures 3E and 3F**). It should be also noted that defective autophagy distinctly affects mitochondrial reorganization in these bryophytes; the mutation in Pp*ATG5* does not affect the total area of mitochondria in spermatids in *P. patens* (Sanchez-Vera et al., 2017), distinct from the corresponding mutation in *M. polymorpha*. Furthermore, whereas autophagy is essential for cytoplasm removal in *M. polymorpha* and *P. patens*, the effects of *atg5* or *atg7* mutations on cytoplasmic removal during spermatozoa formation in mice are less remarkable (Huang et al., 2021; Sanchez-Vera et al., 2017; Shang et al., 2016; Wang et al., 2014). This is not surprising, considering that the cytoplasm should be removed in distinct ways between plants and mammals; the excess cytoplasm is removed by phagocytosis by neighboring Sertoli cells in mammals (O’Donnell et al., 2011), whereas cytoplasm elimination must be accomplished in a cell-autonomous manner in plants because phagocytic removal of the cytoplasm by neighboring cells cannot occur due to the rigid cell wall in plants.

Thus, we demonstrated that multiple degradation mechanisms, including distinct modes of autophagy, play essential roles in male gametogenesis in *M. polymorpha*. Our findings also demonstrated that the mechanisms and roles of degradative pathways underpinning male gametogenesis vary among eukaryotic lineages, even among bryophytes, as shown by the distinct involvement of autophagy in microtubule organization and mitochondrial reorganization between *M. polymorpha* and *P. patens*. The highly dynamic cellular reorganization during bryophyte spermiogenesis would provide a suitable platform for studying various mechanisms of degradative regulation of cellular components in plants, which include selective and nonselective autophagy, endocytic degradation involving ESCRT, and other degradative mechanisms to be discovered in future studies.

## ACKNOWLEDGMENTS

We thank Dr. Takayuki Kohchi (Kyoto University), Dr. Ryuichi Nishihama (Tokyo University of Science), Dr. Shin-ichi Arimura (The University of Tokyo), Dr. Shoji Mano and Dr. Takehiko Kanazawa (National Institute for Basic Biology) for sharing vectors, antibodies, or plant materials. We thank the Model Plant Research Facility, NIBB Bioresource Center, for providing plant cultivation facilities. This study was supported by Grants-in-Aid for Scientific Research from the Ministry of Education, Culture, Sports, Science, and Technology of Japan (20K15824 to N.M., 19H05672 to H.T., and 19H05675, 19H05670, and 21H02515 to T.U.), and a Grant-in-Aid from the Japan Society for the Promotion of Science (JSPS) (19J13751 to TN).

## AUTHOR CONTRIBUTIONS

T.N. performed a major part of experiments. N.M. constructed some vectors and performed a TEM experiment. T.N. and T.U. wrote the manuscript, and H.T. and T.U. supervised the study.

## STAR METHODS

### RESOURCE AVAILABILITY

#### Lead contact

Further information and requests for resources and reagents should be directed to and will be fulfilled by the Lead Contact, Takashi Ueda.

#### Materials availability

Transgenic plants and plasmids generated in this study will be made available on request. Please note that their transfer will be subject to a Materials Transfer Agreement and any relevant import permits.

## Data and code availability

No datasets or code were generated in this study.

## EXPERIMENTAL MODEL AND SUBJECT DETAILS

### Plant materials and growth conditions

The *M. polymorpha* accessions Takaragaike-1 (Tak-1, male) and Takaragaike-2 (Tak-2, female) (Ishizaki et al., 2008) were grown asexually on 1/2× Gamborg’s B5 medium containing 1.4% (w/v) agar at 22 °C under continuous white light. For the induction of sexual reproduction, approximately two-week-old thalli grown on 1/2× Gamborg’s B5 medium containing 1.4% (w/v) agar at 22 °C under continuous white light were further cultivated on vermiculite soaked in 1:1000 Hyponex (HYPONeX JAPAN) at 22 °C under continuous white light for two weeks and then cultivated at 22 °C under continuous white light supplemented with far-red light.

## METHOD DETAILS

### Vector construction

The coding sequence (CDS) and genomic sequence of genes in *M. polymorpha* were amplified by polymerase chain reaction (PCR) from cDNA or genomic DNA prepared from Tak-1 thalli. For the nomenclature of genes, proteins, and mutants of *M. polymorpha*, we followed Bowman et al., (2016). Gene IDs were obtained from MarpolBase (http://marchantia.info/), genome version 5.1 (Montgomery et al., 2020). The primer sequences used in this study are listed in **Table S1**.

For the construction of pENTR *Mito*, cDNA for the amino-terminal 64 amino acids of the gamma subunit of F1ATPase in *M. polymorpha* (Mp7g04400.1) was amplified and subcloned into pENTR/D-TOPO (Thermo Fisher Scientific) according to the manufacturer’s instructions.

For the construction of pENTR *_pro_*Mp*SYP2:Mito*, the 5’ sequence [promoter + 5’ untranslated region (UTR)] of Mp*SYP2* (Kanazawa et al., 2016) was inserted into the *Not*I site of pENTR *Mito* using the In-Fusion HD Cloning System (Clontech).

For the construction of pENTR Mp*IDH1*, the CDS for Mp*IDH1* (Mp1g01980.1) was amplified and subcloned into pENTR/D-TOPO.

For the construction of pENTR *_pro_*Mp*DRP3:3×HA-*Mp*DRP3l*, the *3×HA* sequence was amplified by PCR and subcloned into pENTR/D-TOPO. Then, the 3.4 kb 5’ sequence (promoter + 5’ UTR) of Mp*DRP3* was amplified by PCR and inserted into the *Not*I site of pENTR *3×HA* using the In-Fusion HD Cloning System (pENTR *_pro_*Mp*DRP3:3×HA*). The CDS for MpDRP3l was amplified from pENTR Mp*DRP3l* (Nagaoka et al., 2017) and inserted into the *Asc*I site of pENTR *_pro_*Mp*DRP3:3×HA* using the In-Fusion HD Cloning System.

For the construction of pENTR *mTurquoise2-*Mp*DRP3l^WT^* or pENTR *mTurquoise2-* Mp*DRP3l^K75A^*, a point mutation (75^th^ Lys to Ala) was introduced by PCR using pENTR Mp*DRP3l* (Nagaoka et al., 2017) to generate pENTR Mp*DRP3l^K75A^*. The CDS for *mTurquoise2* was inserted into the *Not*I site of pENTR Mp*DRP3l^WT^* or pENTR Mp*DRP3l^K75A^* using the In-Fusion HD Cloning System.

For the construction of pENTR *_pro_*Mp*ATG5:*Mp*ATG5^res^*, the CDS for Mp*ATG5* (Mp1g12840.1), including the stop codon, was amplified and subcloned into pENTR/D-TOPO. Point mutations (AGACCT to AGGCCA in ^111^Lys-^112^Pro) were introduced into pENTR Mp*ATG5* (+stop codon) by PCR. The 5.0 kb 5’ sequence (promoter + 5’ UTR) of Mp*ATG5* was amplified and inserted into the *Not*I site of this vector using the In-Fusion HD Cloning System.

For the construction of pENTR *_pro_*Mp*ATG5:*Mp*ATG5-mCitrine*, the CDS for MpATG5 without the stop codon was amplified and subcloned into pENTR/D-TOPO. The CDS for *mCitrine* was amplified and inserted into the *Asc*I site of pENTR Mp*ATG5* using the In-Fusion HD Cloning System. The 5.0 kb 5’ sequence (promoter + 5’ UTR) of Mp*ATG5* was amplified and inserted into the *Not*I site of this vector using the In-Fusion HD Cloning System.

For the construction of pENTR *_pro_*Mp*VAMP71:mGFP*-Mp*VAMP71*, the cDNA for *mGFP* was inserted into the *Sma*I site of pENTR *_pro_*Mp*VAMP71*-Mp*VAMP71* previously reported in Minamino et al. (2017) using the In-Fusion HD Cloning System.

For the construction of pENTR *_pro_*Mp*ATG8a:mTurquoise2-*Mp*ATG8a*, the CDS for *mTurquoise2* containing the *Sma*I site at the 5’ end was subcloned into pENTR/D-TOPO. The 5.0 kb 5’ sequence (promoter + 5’ UTR) of Mp*ATG8a* was then amplified and inserted into the *Sma*I site of pENTR *mTurquoise2*. The genomic sequences of Mp*ATG8a* (Mp1g21590.1) comprising the protein-coding regions and 3’ flanking sequences (2.0 kb) were amplified and inserted into the *Asc*I site of pENTR *_pro_*Mp*ATG8a:mTurquoise2* using the In-Fusion HD Cloning System.

For the construction of pENTR *mTurquoise2-PTS1*, the pENTR backbone and the CDS for *mTurquoise2* containing the PTS1 sequence (Val-Ser-Lys-Leu) at the 3’ end were amplified and fused using the In-Fusion HD Cloning System.

For the construction of pENTR *mTurquoise2-*Mp*GOS11*, the CDS for *mTurquoise2* was inserted into the *Not*I site of pENTR Mp*GOS11* (Kanazawa et al., 2016) using the In-Fusion HD Cloning System.

For the construction of pENTR *ST-mTurquoise2*, the sequence for the transmembrane domain of rat ST was amplified from pENTR *ST*-*Venus* (Uemura et al., 2012) and subcloned into pENTR/D-TOPO. The CDS for *mTurquoise2* was inserted into the *Asc*I site of pENTR *ST* using the In-Fusion HD Cloning System.

For the construction of the CRISPR/Cas9 vector, two complementary oligonucleotides in the sequences of Mp*ATG13* and Mp*ATG14* were synthesized and annealed, and the resulting double-stranded fragments were subcloned at the *Bsa*I site of the pMpGE_En03 vector (Sugano et al., 2018) using the DNA ligation kit Ver.2.1 (Takara Bio) according to the manufacturer’s instructions.

For the construction of pMpGWB101 *_pro_*Mp*MS1*, a promoter sequence with a 5.1 kb 5’ sequence (promoter + 5’ UTR) of Mp*MS1* (Mp3g17000.1) was introduced at the *Hin*dIII site of pMpGWB101 (Ishizaki et al., 2015) using the In-Fusion HD Cloning System.

For the construction of pMpGWB301 *_pro_*Mp*SYP2:*Gateway-*mCitrine*, the amplicons of *_pro_*Mp*SYP2* (Kanazawa et al., 2016) and cDNA for *mCitrine* were inserted into the *Hin*dIII and *Sac*I sites of pMpGWB301 (Ishizaki et al., 2015), respectively, using the In-Fusion HD Cloning System.

The sequences flanked by the *att*L1 and *att*L2 sites in the resultant entry vectors were introduced into the destination vectors using Gateway LR Clonase™ II Enzyme Mix (Thermo Fisher Scientific) according to the manufacturer’s instructions, as listed in **Table S2**. The pMpGWB301 *_pro_*Mp*DUO1*:*mCitrine*-Mp*SEC22*, pMpGWB301 *_pro_*Mp*DUO1*:*mCitrine*-Mp*GOS11*, pMpGWB301 *_pro_*Mp*DUO1*:*ST*-*Venus*, pMpGWB303 *Citrine-PTS1*, pMpGWB301 *_pro_*Mp*SYP12A*:*mCitrine*-Mp*SYP12A*, and pMpGWB301 *_pro_*Mp*VAMP71*:*mCitrine*-Mp*VAMP71* vectors, which were previously constructed (Kanazawa et al., 2016; Mano et al., 2018; Minamino et al., 2017; Minamino et al., 2021) were also used in this study.

### Domain search

A domain search of MpATG13 and MpATG14 was performed using SMART (http://smart.embl-heidelberg.de/) (Letunic and Bork, 2018; Letunic et al., 2021).

### Transformation and crossing

The transformation of *M. polymorpha* was performed as previously described (Kubota et al., 2013). Two-week-old Tak-1 thalli, whose apical parts, including meristems, were removed, were grown on 1/2× Gamborg’s B5 medium containing 1.0% (w/v) sucrose and 1.4% (w/v) agar at 22 °C under continuous white light for four days to regenerate the thalli. The regenerating plantlets were cocultivated with the agrobacterium strain GV2260 harboring the binary vector in liquid 0M51C medium containing 2% (w/v) sucrose and 100 μM acetosyringone (Sigma-Aldrich) at 22 °C under continuous white light for four days. Transformants were selected on plates containing 10 mg/L hygromycin B and 250 mg/L cefotaxime for the pMpGWB101, pMpGWB103, and pMpGE010 vectors and 0.5 µM chlorsulfuron and 250 mg/L cefotaxime for the pMpGWB301, pMpGWB303, and pMpGWB308 vectors.

To cross male and female lines, spermatozoids were obtained by placing a mature antheridiophore upside-down on a drop of 20–30 μL of water for one minute and then applied to archegoniophores. Archegoniophores with yellow sporangia were obtained between three and five weeks after crossing.

### Genotyping

For the genotyping of mutants generated by the CRISPR/Cas9 system, total RNA was extracted from 5-day-old wild-type and Mp*atg* mutant thalli using the RNeasy Plant Mini Kit (Qiagen) and used as a template for reverse transcription using SuperScript III Reverse Transcriptase (Thermo Fisher Scientific) and the oligo (dT) (18-mer) primer according to the manufacturer’s instructions. Mutations in the obtained cDNA fragments were analyzed by direct sequencing.

For the generation of the male Mp*atg5-1^ge/cf^* plant, crossing of Mp*atg5-1^ge^*/*_pro_*Mp*ATG5*:Mp*ATG5*^res^ (male) with Tak-2 (female) was performed (**Figure S6**). The obtained F1 progeny were grown on 1/2× Gamborg’s B5 medium containing 10 mg/L hygromycin and 250 mg/L cefotaxime or 0.5 µM chlorsulfuron and 250 mg/L cefotaxime, and F1 progenies harboring neither hygromycin-nor chlorsulfuron-resistant genes were isolated. To check whether these progeny were male plants with the Mp*atg5-1^ge^* mutation, genomic DNA was extracted from the thalli in genotyping buffer containing 1 M KCl, 100 mM Tris-HCl (pH 9.5), and 10 mM EDTA and used as the template for PCR using KOD FX Neo (TOYOBO) according to the manufacturer’s instructions. The sex of the obtained plants was checked by PCR using the primers listed in **Table S1** as previously described (Fujisawa et al., 2001). Mutations in the obtained genomic DNA fragments were analyzed by direct sequencing.

### Cell wall digestion of antheridial cells

Cell wall digestion was carried out according to Shimamura (2015). Antheridia were fixed with 4% (w/v) paraformaldehyde (PFA) in PME buffer [50 mM PIPES, 5 mM EGTA, and 1 mM MgSO_4_, adjusted at pH 6.8 using NaOH] for 60 minutes and then washed with PME buffer three times. The fixed antheridia were incubated in wall-digesting enzyme solution [1% (w/v) Cellulase Onozuka RS (SERVA), 0.25% (w/v) Pectolyase Y-23 (Kyowa Chemical Products), 1% (w/v) bovine serum albumin (BSA), 0.1% (w/v) IGEPAL CA-630, 1× cOmplete^TM^ protease inhibitor cocktail (Roche), and 1% (w/v) glucose in PME buffer] for 30 minutes and then washed with PME buffer three times. The cell wall-digested antheridia were placed on MAS-coated glass slides (Matsunami Glass), and a coverslip was placed on the sample and pressed down gently with a finger to squash the sample. The coverslip was removed, and the sample was incubated in PME buffer containing 1 µg/mL Hoechst 33342 solution or propidium iodide.

### Immunostaining of antheridial cells

Cell wall digestion was performed as described above, and immunostaining was carried out according to Shimamura (2015). After incubation with the wall-digesting enzyme solution, the sample was incubated in permeabilization buffer [1% (w/v) BSA and 0.1 or 0.01% (v/v) Triton X-100 in PME buffer] for 10 minutes and then washed with PME buffer three times. The cell wall-digested antheridia were placed on MAS-coated glass slides, and a coverslip was placed on the sample and pressed gently with a finger to squash the sample. After the coverslip was removed, the sample was incubated in 1% (w/v) BSA in PBS buffer [150 mM NaCl, 80 mM Na_2_HPO_4_, and 40 mM NaH_2_PO_4_ (adjusted at pH 6.8 using NaOH)] for 30 minutes and then incubated in primary antibody solution [1:200 for the anti-HA antibody (M180-3; Medical & Biological Laboratories), 1:1000 for the anti-α-tubulin antibody (DM1A; Sigma-Aldrich), or 1:1000 for the anti-catalase antibody (Yamaguchi and Nishimura, 1984) in PBS buffer containing 1% (w/v) BSA] overnight at 4 °C. The samples were washed with PBS buffer three times and incubated in secondary antibody solution [1:1000 Alexa Fluor™ 546 goat anti-mouse IgG (H+L) (Thermo Fisher Scientific) for use in experiments with anti-HA and anti-catalase antibodies as the primary antibodies and 1:1000 Alexa Fluor™ 488 goat anti-mouse IgG (H+L) (Thermo Fisher Scientific) for use in experiments with the anti-α-tubulin antibody as the primary antibody, both in PBS buffer containing 1% (w/v) BSA] for at least one hour at 37 °C. The samples were washed with PBS buffer three times and incubated in 1 µg/mL Hoechst 33342 in PBS buffer for at least 10 minutes. The samples were mounted with ProLong™ Diamond Antifade Mountant (Thermo Fisher Scientific) and incubated for at least 24 hours at room temperature in the dark.

### Confocal laser scanning microscopy

For the observation of the mitochondrial marker in thallus cells, five-day-old thalli expressing each of the markers were stained with 0.5 M MitoTracker^TM^ Orange CMTMRos (Thermo Fisher Scientific) for 20 minutes and washed three times with 1/2× Gamborg’s B5 liquid medium, as previously described (Nagaoka et al., 2017). For the observation of organelle markers in antheridial cells, antheridial cells whose cell wall was digested as described above or antheridia hand-sectioned manually by a razor blade were used. For the observation of spermatozoids, spermatozoids were obtained as described above and fixed in 4% (w/v) PFA in PBS buffer for five minutes. Fixed spermatozoids were centrifuged at 9,100 ×g for one minute, and the pellet was suspended in PBS buffer with or without 1 µg/mL Hoechst 33342.

For confocal microscopic observation, an LSM780 confocal microscope (Carl Zeiss) equipped with an oil immersion lens (×63, numerical aperture = 1.4) was used.

For measurement of the total area of mitochondria per cell, maximum-intensity projection images created from z-stacked images by ImageJ (version 1.50i) were used, and the mitochondrial area was measured by ImageJ. The circularity of nuclei was calculated using ImageJ from single confocal images.

### Dark field microscopy

To trace the movement of spermatozoids, spermatozoids collected as described above were observed with a dark-field microscope (Olympus) equipped with an ORCA-Flash4.0 V2 camera (Hamamatsu photonics). To obtain the trajectories of spermatozoids, 33 frames taken every 30 milliseconds were processed with the ImageJ macro, Color Footprint Rainbow.

### Electron microscopy

Wild-type and Mp*atg5-1^ge^* antheridia were subjected to electron microscopic observation. Sample preparation was performed according to Minamino et al. (2017). The antheridia were fixed with 2% (w/v) PFA and 2% (v/v) glutaraldehyde in 0.05 M cacodylate buffer (pH 7.4) at 4 °C overnight. The fixed samples were washed three times with 0.05 M cacodylate buffer for 30 minutes each and were then postfixed with 2% (w/v) osmium tetroxide in 0.05 M cacodylate buffer at 4 °C for three hours. The samples were dehydrated in graded ethanol solutions [50 and 70% (v/v) ethanol for 30 minutes each at 4 °C, 90% (v/v) for 30 minutes at room temperature, four times with 100% for 30 minutes each at room temperature, and 100% overnight at room temperature]. The samples were infiltrated with propylene oxide (PO) two times for 30 minutes each and then placed into a 70:30 mixture of PO and resin (Quetol-651; Nisshin EM) for one hour. The caps of tubes were opened overnight to volatilize the PO. The samples were transferred to fresh 100% resin and polymerized at 60 °C for 48 hours. Ultrathin sample sections were mounted on copper grids, stained with 2% (w/v) uranyl acetate and lead stain solution (Sigma-Aldrich), and observed with a transmission electron microscope (JEM-1400Plus; JEOL) at an acceleration voltage of 80 kV. Digital images (3296 × 2472 pixels) were taken with a CCD camera (EM-14830RUBY2; JEOL).

### Immunoblot analysis

Immunoblot analysis was carried out as described in Norizuki et al. (2019). Gemma were grown on cellophane placed on 1/2× Gamborg’s B5 agar medium for five days. 50 mg of plants of each genotype were homogenized in 100 µL of grinding buffer [50 mM HEPES– KOH (pH 7.5), 340 mM sorbitol, 5 mM MgCl_2_, and 1× cOmplete^TM^ Protease Inhibitor Cocktail] and centrifuged at 1,000× g for 10 minutes. The supernatants were centrifuged at 3,000× g for 10 minutes, and the resulting supernatants were used for immunoblotting. The polyclonal anti-GFP antibody purified by affinity column chromatography using the GST-mCitrine protein (Norizuki et al., 2019) was used at ×1,000 dilution as the primary antibody. The peroxidase-conjugated donkey anti-rabbit immunoglobulin antibody (GE Healthcare) was used as the secondary antibody. Signals were detected using Immobilon^TM^ Western Chemiluminescent HRP Substrate (Merck Millipore).

## QUANTIFICATION AND STATISTICAL ANALYSIS

To test the normality of the data, the Shapiro-Wilk test was performed using R (version 3.6.0), and samples were considered nonparametric when the *p* value was less than 0.05. For comparison between two groups, the Welch’s t-test (for parametric samples) or Wilcoxon rank-sum test (for nonparametric samples) was performed using the software R. For statistical analyses among three or more groups, the Steel–Dwass test was performed using JMP 16 (SAS Institute). Statistical method details are indicated in each figure legend.

## SUPPLEMENTAL INFORMATION

Figure S1. Reorganization of mitochondria during spermiogenesis in *M. polymorpha*, related to Figure 1 (A and B) Confocal images of thallus cells expressing *_pro_*Mp*EF1α:Mito*-*Citrine* (A) or *_pro_*Mp*SYP2:*Mp*IDH1*-*mCitrine* (B) (green) and stained with MitoTracker^TM^ Orange CMTMRos (MitoTracker; magenta). MitoTracker labeled the mitochondria and the apoplast. The lower panels are enlarged images of the regions enclosed in squares. Blue pseudocolor indicates the fluorescence from autofluorescence from chlorophyll. Scale bars = 20 μm. (C) Maximum-intensity projection images of cell wall-digested antheridial cells and a spermatozoid expressing *_pro_*Mp*SYP2:*Mp*IDH1*-*mCitrine* (green). Nuclei were stained with Hoechst 33342 (blue). Arrowheads indicate the elongated mitochondria located beneath the base of flagella. Scale bars = 5 μm. (D) Total area of mitochondria in cells at stages 0 to 5 calculated using maximum-intensity projection images including samples presented in (C). n = 52 (stage 0), 51 (stage 1), 54 (stage 2), 56 (stage 3), and 30 (stages 4 and 5) cells. The boxes and solid lines in the boxes indicate the first and third quartile and the median, respectively. The whiskers indicate 1.5× interquartile ranges. Different letters denote significant differences based on the Steel– Dwass test (*p* < 0.05). (E) The number of mitochondria counted in the same sets of samples analyzed in (D). The bold letter is the most frequent mitochondrial number in each stage. n: the number of cells.

Figure S2. Dominant negative effect of MpDRP3l^K75A^ on the number of mitochondria in spermatids, related to Figure 2 (A) Part of the amino acid sequence alignment of ScDnm1, AtDRP3B, and MpDRP3. The red letters show the lysine residue, which was replaced with alanine to construct the dominant-negative form of dynamin. Sc: *Saccharomyces cerevisiae*, At: *Arabidopsis thaliana*. (B) Maximum-intensity projection images of stage 1 cells expressing *_pro_*Mp*EF1α:Mito-Citrine* (green) alone (left two panels) or with *_pro_*Mp*MS1:mTurquoise2-*Mp*DRP3l ^K75A^* (magenta, middle two panels) or *_pro_*Mp*MS1:mTurquoise2-*Mp*DRP3l^WT^* (magenta, right two panels). Nuclei were stained with propidium iodide (PI; blue). Scale bars = 5 μm. (C) The percentage of stage 1 cells with only one mitochondrion, which was calculated using samples including those presented in (B). n: the number of cells. KA1–3: three independent lines of spermatozoids expressing mTurquoise2-MpDRP3l^K75A^. WT1, 4, and 5: three independent lines of spermatozoids expressing mTurquoise2-MpDRP3l^WT^. (D) Spermatozoids with the indicated types of mitochondria expressing *_pro_*Mp*EF1α:Mito*-*Citrine* (green) and *_pro_*Mp*MS1:mTurquoise2-*Mp*DRP3l^K75A^* (not shown). The arrow shows the posterior mitochondrion, and the arrowheads show additional mitochondria observed between the anterior and posterior mitochondria. Scale bars = 5 μm. (E) Percentage of each type of spermatozoid expressing *_pro_*Mp*EF1α:Mito*-*Citrine* and *_pro_*Mp*MS1:mTurquoise2-*Mp*DRP3l^WT^* (WT) or *_pro_*Mp*MS1:mTurquoise2-*Mp*DRP3l^K75A^* (KA). n: the number of spermatozoids. (F) Total area of mitochondria in spermatozoids expressing *_pro_*Mp*EF1α:Mito*-*Citrine* and *_pro_*Mp*MS1:mTurquoise2-*Mp*DRP3l^WT^* or *_pro_*Mp*MS1:mTurquoise2-*Mp*DRP3l^K75A^*. Mitochondrial area was calculated from maximum-intensity projection images. n = 34 (KA1), 35 (KA2 and KA3), 20 (WT1), and 30 (WT4 and WT5) spermatozoids. Different letters denote significant differences based on the Steel–Dwass test (*p* < 0.05).

Figure S3. Mp*atg13* and Mp*atg14* mutants are defective in autophagy and spermiogenesis, related to Figure 3 (A and B) Gene models of Mp*ATG13* (A) and Mp*ATG14* (B). The black and gray boxes indicate the CDS and UTR, respectively. gRNAs indicate positions of gRNA used for generating mutants with the CRISPR/Cas9 system. gRNA1 and gRNA2 were used to generate Mp*atg14-1^ge^* and Mp*atg14-2^ge^*, respectively. (C and D) Domain structures of MpATG13 (C) and MpATG14 (D). (E and F) Sequences of the mutations in Mp*ATG13* (E) and Mp*ATG14* (F) introduced by genome editing. Underlines indicate the PAM sequences. Inserted and deleted bases and substituted amino acid residues are shown in red. The asterisk indicates the stop. (G) Five-day-old thalli of Mp*atg13^ge^* (left) and Mp*atg14^ge^* (right) expressing mCitrine-MpATG8a were subjected to immunoblotting using the anti-GFP antibody. Wild type and Mp*atg5-1*^ge^ expressing mCitrine-MpATG8a were also included as positive and negative controls. The amount of free mCitrine was drastically reduced in Mp*atg13^ge^* and Mp*atg14^ge^*, indicating impaired transport of mCitrine-MpATG8a in these mutants. (H) Confocal images of Mp*atg13-2^ge^* and Mp*atg14-2^ge^* spermatozoids. Nuclei were stained with Hoechst 33342 (blue). Scale bars = 5 μm.

Figure S4. Male sterility of Mp*atg* mutants, related to Figure 3 (A) Male wild-type or Mp*atg* mutant plants were crossed with female wild-type plants, and archegoniophores were observed after three to five weeks. Arrowheads indicate yellow sporangia. Bars = 0.5 cm. (B) The number of archegoniophores with yellow sporangia after crossing.

Figure S5. Impaired reorganization of mitochondria during spermiogenesis in autophagy-defective mutants, related to Figure 4 (A) Confocal images of wild-type and Mp*atg5-1^ge^* spermatozoids expressing *_pro_*Mp*SYP2:*Mp*IDH1*-*mCitrine* (green). Nuclei were stained with Hoechst 33342 (blue). Scale bars = 5 μm. (B) The total area of mitochondria in wild-type and Mp*atg5-1^ge^* spermatozoids. The mitochondrial area was calculated in maximum-intensity projection images of samples including spermatozoids presented in (A). n = 22 (wild-type) and 24 (Mp*atg5-1^ge^*) spermatozoids. The boxes and solid lines in the boxes indicate the first quartile and third quartile and the median, respectively. The whiskers indicate 1.5× interquartile ranges. Statistical differences were analyzed by the Welch’s t-test. (C) Maximum-intensity projection images of stages 0, 1, and 3 cells in Mp*atg5-1^ge^* expressing *_pro_*Mp*EF1α:Mito*-*Citrine* (green). Nuclei were stained with Hoechst 33342 (blue). Arrowheads indicate the elongated mitochondria. Scale bars = 5 μm.

Figure S6. Generation of the Mp*atg5* mutant without the T-DNA cassette comprising *CAS9*, related to Figure 6 and STAR Methods (A) The T-DNA cassette used for generating the Mp*atg5-1^ge^* mutant. (B) Confocal images of wild-type, Mp*atg5-1^ge/cf^*, and Mp*atg5-1^ge/cf^* spermatozoids expressing *_pro_*Mp*ATG5:*Mp*ATG5-mCitrine*. Nuclei were stained with Hoechst 33342 (blue). Scale bars = 5 μm.

Table S1. Primer list, related to STAR Methods

Table S2. **List of vectors used in LR recombination, related to STAR Methods** S1: Ishizaki et al. (2015), S2; Minamino et al., (2021), S3; Sugano et al., (2018).

