## Supplementary material for "Bryophyte spermiogenesis occurs through multimode autophagic and nonautophagic degradation": Supplemenental Figures

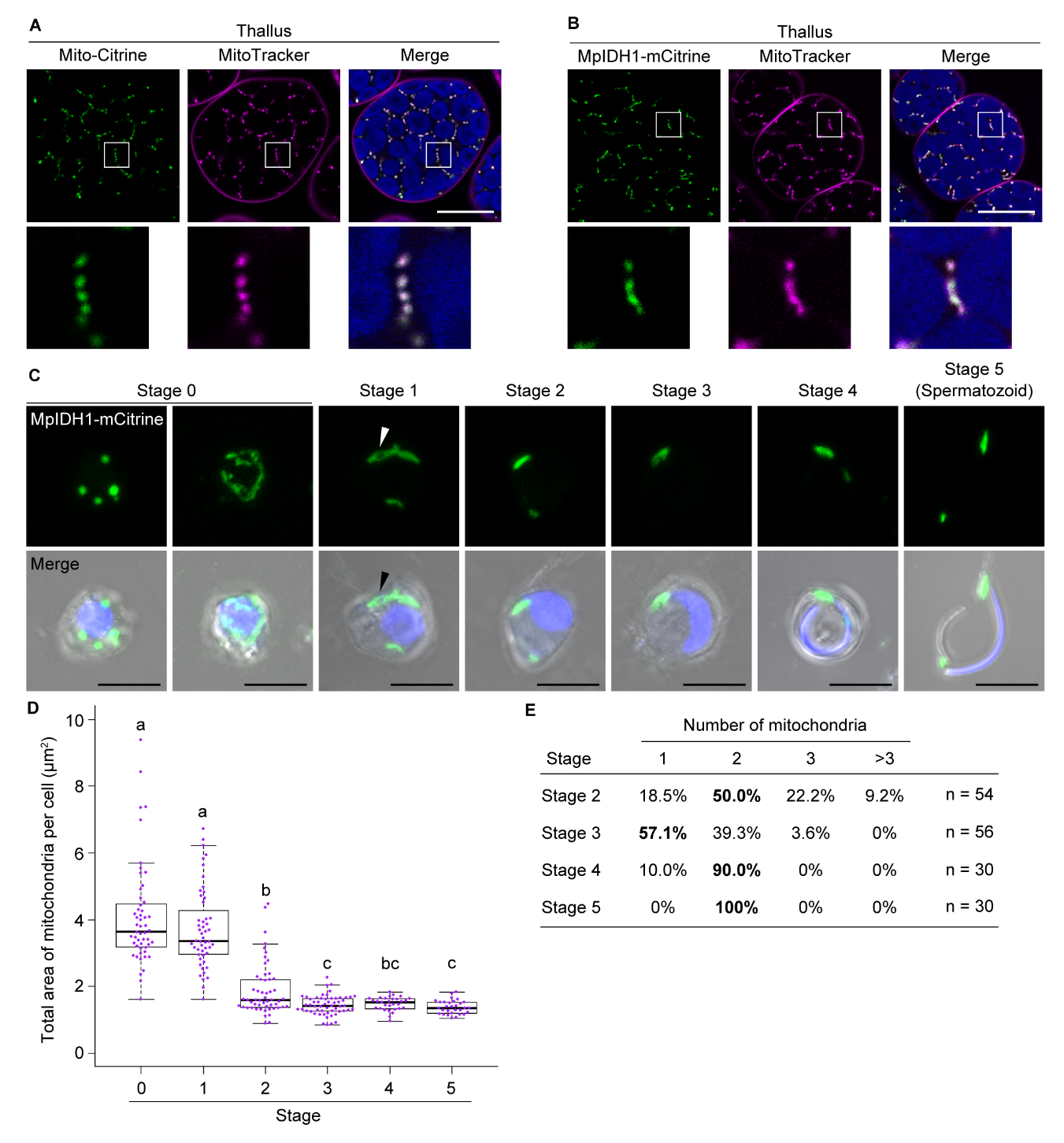

**Figure S1. Reorganization of mitochondria during spermiogenesis in *M. polymorpha*, related to Figure 1**

(A and B) Confocal images of thallus cells expressing *_pro_*Mp*EF1α:Mito*-*Citrine* (A) or *_pro_*Mp*SYP2:*Mp*IDH1*-*mCitrine* (B) (green) and stained with MitoTracker^TM^ Orange CMTMRos (MitoTracker; magenta). MitoTracker labeled the mitochondria and the apoplast. The lower panels are enlarged images of the regions enclosed in squares. Blue pseudocolor indicates the fluorescence from autofluorescence from chlorophyll. Scale bars = 20 μm.

(C) Maximum-intensity projection images of cell wall-digested antheridial cells and a spermatozoid expressing *_pro_*Mp*SYP2:*Mp*IDH1*-*mCitrine* (green). Nuclei were stained with Hoechst 33342 (blue). Arrowheads indicate the elongated mitochondria located beneath the base of flagella. Scale bars = 5 μm.

(D) Total area of mitochondria in cells at stages 0 to 5 calculated using maximum-intensity projection images including samples presented in (C). n = 52 (stage 0), 51 (stage 1), 54 (stage 2), 56 (stage 3), and 30 (stages 4 and 5) cells. The boxes and solid lines in the boxes indicate the first and third quartile and the median, respectively. The whiskers indicate 1.5× interquartile ranges. Different letters denote significant differences based on the Steel–Dwass test (*p* < 0.05).

(E) The number of mitochondria counted in the same sets of samples analyzed in (D). The bold letter is the most frequent mitochondrial number in each stage. n: the number of cells.

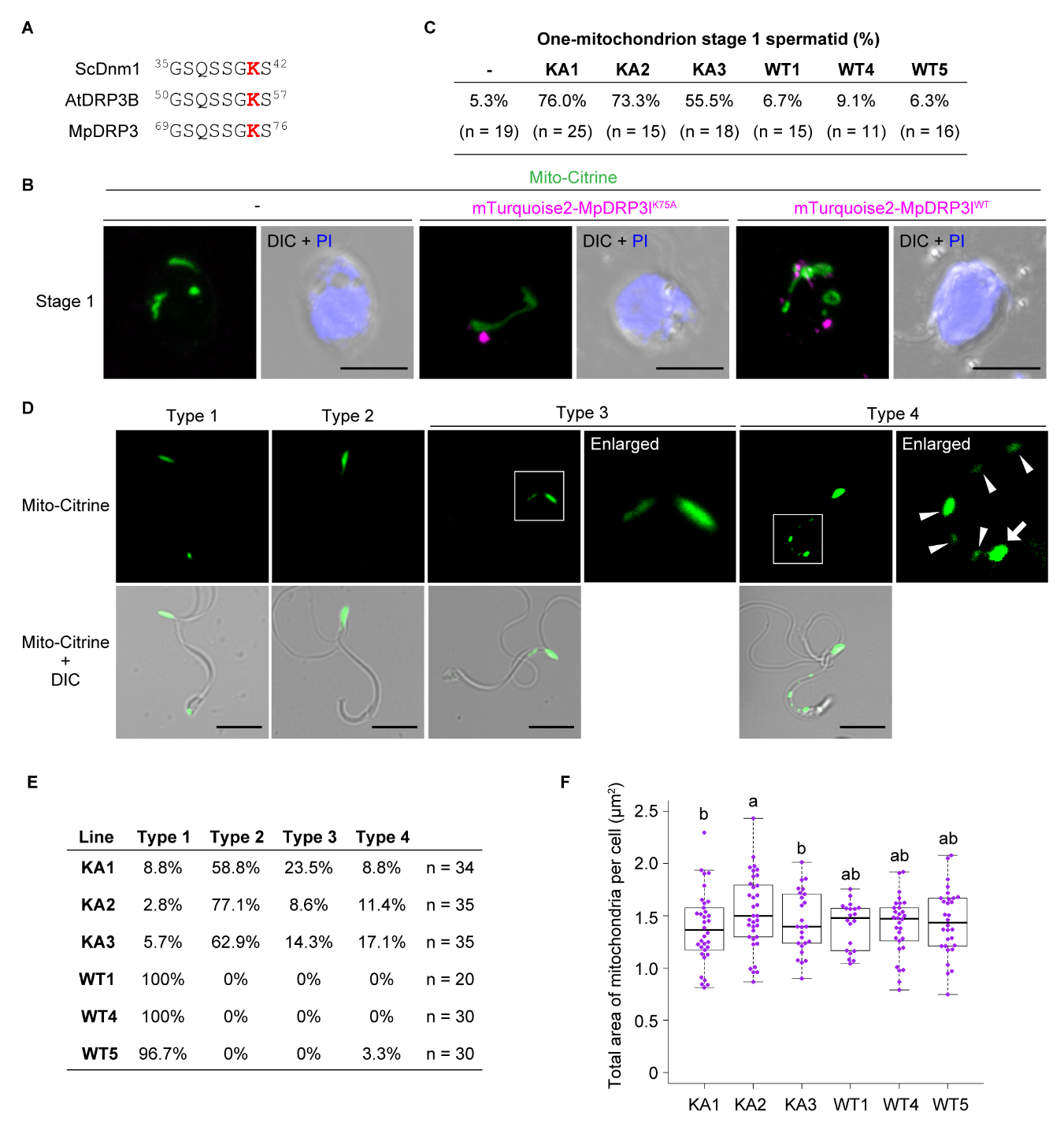

**Figure S2. Dominant negative effect of MpDRP3l^K75A^ on the number of mitochondria in spermatids, related to Figure 2**

(**A**) Part of the amino acid sequence alignment of ScDnm1, AtDRP3B, and MpDRP3. The red letters show the lysine residue, which was replaced with alanine to construct the dominant-negative form of dynamin. Sc: *Saccharomyces cerevisiae*, At: *Arabidopsis thaliana*.

(B) Maximum-intensity projection images of stage 1 cells expressing *_pro_*Mp*EF1α:Mito*-*Citrine* (green) alone (left two panels) or with *_pro_*Mp*MS1:mTurquoise2-*Mp*DRP3l ^K75A^* (magenta, middle two panels) or *_pro_*Mp*MS1:mTurquoise2-*Mp*DRP3l^WT^* (magenta, right two panels). Nuclei were stained with propidium iodide (PI; blue). Scale bars = 5 μm.

(C) The percentage of stage 1 cells with only one mitochondrion, which was calculated using samples including those presented in (B). n: the number of cells. KA1–3: three independent lines of spermatozoids expressing mTurquoise2-MpDRP3l^K75A^. WT1, 4, and 5: three independent lines of spermatozoids expressing mTurquoise2-MpDRP3l^WT^.

(D) Spermatozoids with the indicated types of mitochondria expressing *_pro_*Mp*EF1α:Mito*-*Citrine* (green) and *_pro_*Mp*MS1:mTurquoise2-*Mp*DRP3l^K75A^* (not shown). The arrow shows the posterior mitochondrion, and the arrowheads show additional mitochondria observed between the anterior and posterior mitochondria. Scale bars = 5 μm.

(E) Percentage of each type of spermatozoid expressing *_pro_*Mp*EF1α:Mito*-*Citrine* and *_pro_*Mp*MS1:mTurquoise2-*Mp*DRP3l^WT^* (WT) or *_pro_*Mp*MS1:mTurquoise2-*Mp*DRP3l^K75A^* (KA). n: the number of spermatozoids.

(F) Total area of mitochondria in spermatozoids expressing *_pro_*Mp*EF1α:Mito*-*Citrine* and *_pro_*Mp*MS1:mTurquoise2-*Mp*DRP3l^WT^* or *_pro_*Mp*MS1:mTurquoise2-*Mp*DRP3l^K75A^*. Mitochondrial area was calculated from maximum-intensity projection images. n = 34 (KA1), 35 (KA2 and KA3), 20 (WT1), and 30 (WT4 and WT5) spermatozoids. Different letters denote significant differences based on the Steel–Dwass test (*p* < 0.05).

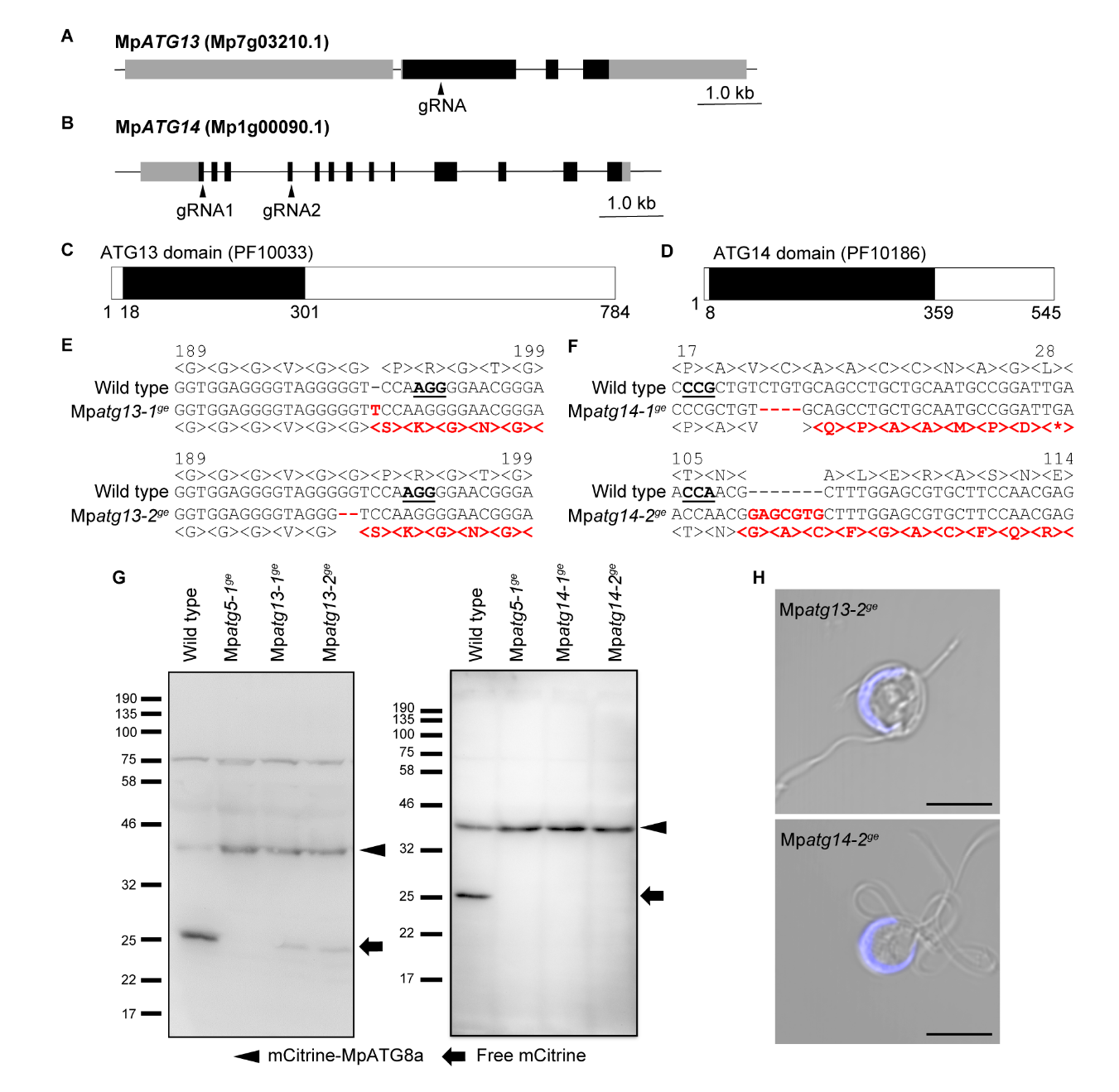

**Figure S3. Mp*atg13* and Mp*atg14* mutants are defective in autophagy and spermiogenesis, related to Figure 3**

(A and B) Gene models of Mp*ATG13* (A) and Mp*ATG14* (B). The black and gray boxes indicate the CDS and UTR, respectively. gRNAs indicate positions of gRNA used for generating mutants with the CRISPR/Cas9 system. gRNA1 and gRNA2 were used to generate Mp*atg14-1^ge^* and Mp*atg14-2^ge^*, respectively.

(C and D) Domain structures of MpATG13 (C) and MpATG14 (D).

(E and F) Sequences of the mutations in Mp*ATG13* (E) and Mp*ATG14* (F) introduced by genome editing. Underlines indicate the PAM sequences. Inserted and deleted bases and substituted amino acid residues are shown in red. The asterisk indicates the stop.

(G) Five-day-old thalli of Mp*atg13^ge^* (left) and Mp*atg14^ge^* (right) expressing mCitrine-MpATG8a were subjected to immunoblotting using the anti-GFP antibody. Wild type and Mp*atg5-1*^ge^ expressing mCitrine-MpATG8a were also included as positive and negative controls. The amount of free mCitrine was drastically reduced in Mp*atg13^ge^* and Mp*atg14^ge^*, indicating impaired transport of mCitrine-MpATG8a in these mutants.

(H) Confocal images of Mp*atg13-2^ge^* and Mp*atg14-2^ge^* spermatozoids. Nuclei were stained with Hoechst 33342 (blue). Scale bars = 5 μm.

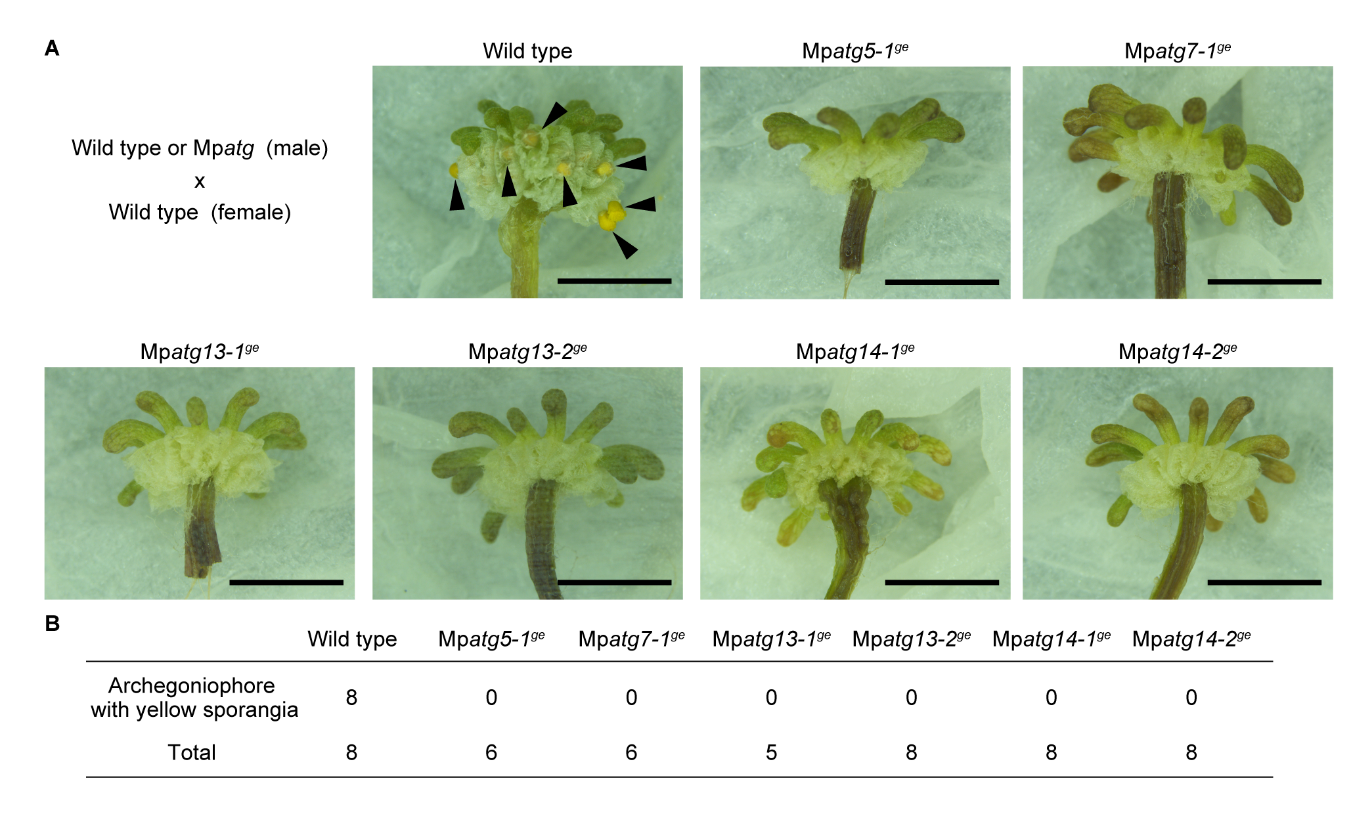

**Figure S4. Male sterility of Mp*atg* mutants, related to Figure 3**

(A) Male wild-type or Mp*atg* mutant plants were crossed with female wild-type plants, and archegoniophores were observed after three to five weeks. Arrowheads indicate yellow sporangia. Bars = 0.5 cm.

(B) The number of archegoniophores with yellow sporangia after crossing.

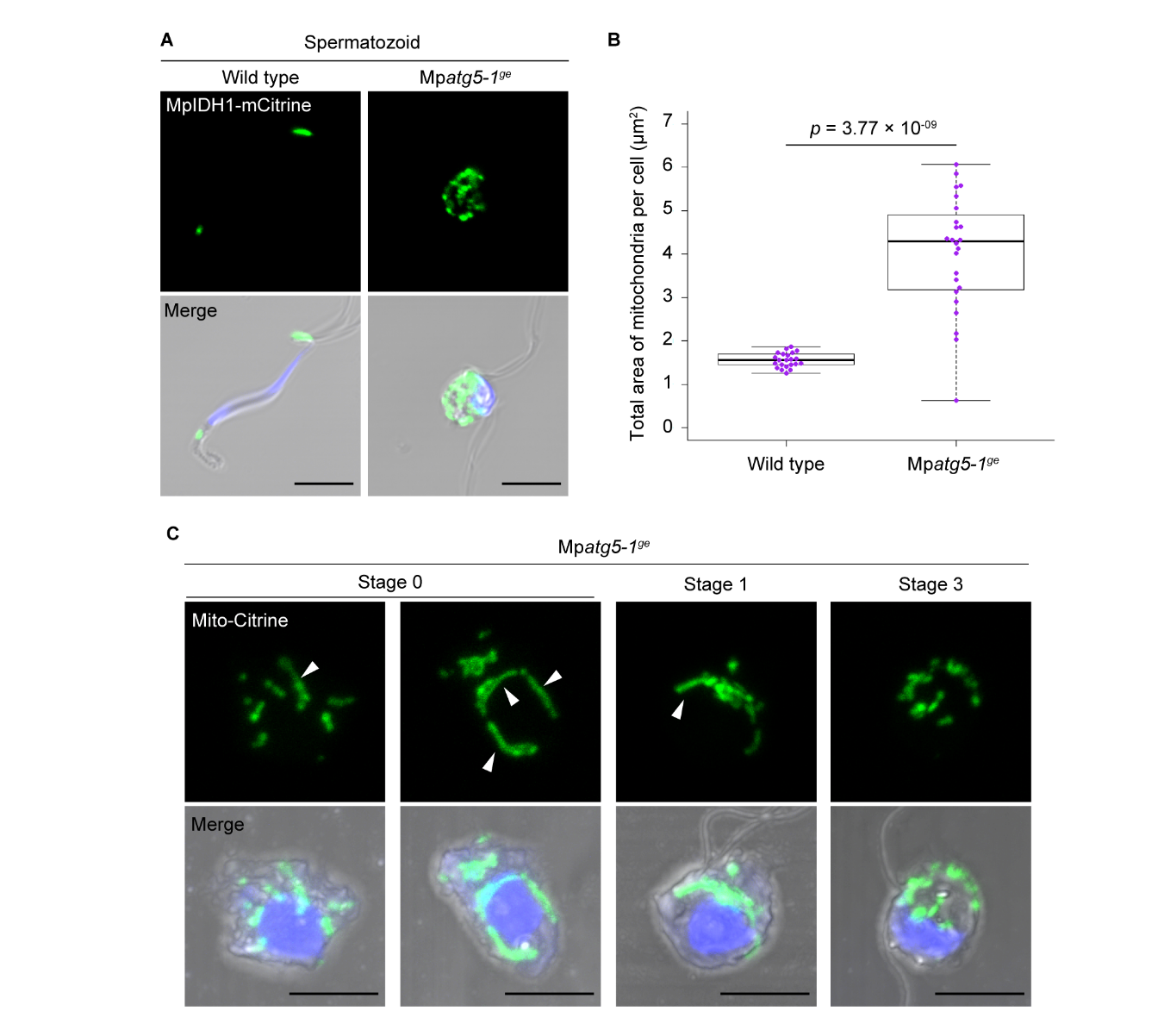

**Figure S5. Impaired reorganization of mitochondria during spermiogenesis in autophagy-defective mutants, related to Figure 4**

(A) Confocal images of wild-type and Mp*atg5-1^ge^* spermatozoids expressing *_pro_*Mp*SYP2:*Mp*IDH1*-*mCitrine* (green). Nuclei were stained with Hoechst 33342 (blue). Scale bars = 5 μm.

(B) The total area of mitochondria in wild-type and Mp*atg5-1^ge^* spermatozoids. The mitochondrial area was calculated in maximum-intensity projection images of samples including spermatozoids presented in (A). n = 22 (wild-type) and 24 (Mp*atg5-1^ge^*) spermatozoids. The boxes and solid lines in the boxes indicate the first quartile and third quartile and the median, respectively. The whiskers indicate 1.5× interquartile ranges. Statistical differences were analyzed by the Welch’s t-test.

(C) Maximum-intensity projection images of stages 0, 1, and 3 cells in Mp*atg5-1^ge^* expressing *_pro_*Mp*EF1α:Mito*-*Citrine* (green). Nuclei were stained with Hoechst 33342 (blue). Arrowheads indicate the elongated mitochondria. Scale bars = 5 μm.

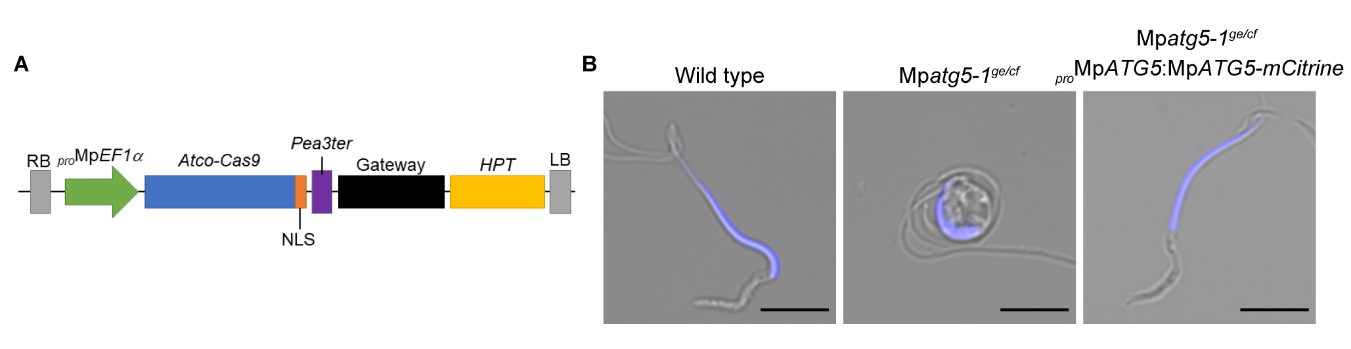

**Figure S6. Generation of the Mp*atg5* mutant without the T-DNA cassette comprising *CAS9*, related to Figure 6 and STAR Methods**

(A) The T-DNA cassette used for generating the Mp*atg5-1^ge^* mutant.

(B) Confocal images of wild-type, Mp*atg5-1^ge/cf^*, and Mp*atg5-1^ge/cf^* spermatozoids expressing *_pro_*Mp*ATG5:*Mp*ATG5-mCitrine*. Nuclei were stained with Hoechst 33342 (blue). Scale bars = 5 μm.

**Table S1. Primer list, related to STAR Methods**

|  | Primer 1 (5´->3') | Primer 2 (5´->3') |
| --- | --- | --- |
| *Mito* | CACCATGGCTTCCGGAAGTGCACG | GGACTTCACAGATTTCATCCTATTTCTC |
| Mp*IDH1* | CACCATGGCTAAGCTCAGGGCTTC | AACATCCAACTTTTCAAGTACAGCA |
| *3×HA* | CACCATGTACCCATACGACGTTCCGGATTACGCTTACCCTTACGACGTGCCGGATTACGC | TCCGCTTCCTCCAGCGTAATCCGGTACGTCGTATGGGTAAGCGTAATCCGGCACGTCGTA |
| *_pro_*Mp*DRP3* | GCAGGCTCCGCGGCCGTTTTCAAATCTATCTCACACGGAGG | GTGAAGGGGGCGGCCTGGGCCAATCGTTGGATGC |
| Mp*DRP3l* | AAGGGTGGGCGCGGGGCGCAGCCTCAAGCAAATC | AGCTGGGTCGGCGCGTCAAAAAGACAAGGTGTCCAGTTC |
| Mp*DRP3l^K75A^* | AGTGGGGCGTCGAGTGTCCTGGAAGCTATGGTGGG | ACTCGACGCCCCACTGCTCTGGCTACCAACGACG |
| *mTurquoise2* (*Not*I) | GCAGGCTCCGCGGCCATGGTGTCTAAGGGTGAGGAACT | GAAGGGGGCGGCCGCTTTGTAAAGCTCATCCATTCCGAGG |
| Mp*ATG5* (+stop) | CACCATGGGGAATGAAGATGAAGATGGA | TCATCTAGAAACCCATGTACATATGCA |
| Mp*ATG5^res^* | TGCTGTGTGCAGAACCGGAGAGGCCATGGAACT | GTTCTGCACACAGCAAGTCGAAAAGGACGCCTGTG |
| *_pro_*Mp*ATG5* | GCAGGCTCCGCGGCCGAAGGAGGTCCTTCCAAAGGCAGAG | GTGAAGGGGGCGGCCCAATATTCACAGCATGAACAAACCT |
| Mp*ATG5*  (-stop) | CACCATGGGGAATGAAGATGAAGATGGA | TCTAGAAACCCATGTACATATGCAAAG |
| *mCitrine* (*Asc*I) | AAGGGTGGGCGCGCCATGGTGAGCAAGGGCGAGGAG | AGCTGGGTCGGCGCGTTACTTGTACAGCTCGTCCATGC |
| *mGFP* (*Sma*I) | GCCCCCTTCACCCCCATGGTGAGCAAGGGCGAGGAG | CATGCCGCTGCCCCCCTTGTACAGCTCGTCCATGCC |
| *mTurquoise2* (pENTR) | CCCTTCACCCCCGGGATGGTGTCTAAGGGTGAGGAACT | GGCGCGCCCACCCTTTTTGTAAAGCTCATCCATTCCGAGG |
| pENTR | AAGGGTGGGCGCGCCGACC | GGTGAAGGGGGCGGCCG |
| *_pro_*Mp*ATG8a* | GCCCCCTTCACCCCCAGGTCTAGTAATCGCTCACTTAGGG | CTTAGACACCATCCCCTTGCCGCTGCTACTACTTTCTTCT |
| Mp*ATG8a* | AAGGGTGGGCGCGGGATGACGGGGAAGAGGAGTTCGTTCA | AGCTGGGTCGGCGCGCTGACAAAGTATCAGTCGTAGGCTC |
| *ST* | CACCATGATTCATACCAACTTGAAGAAAAAGTTC | CATGGCCACTTTCTCCTGGC |
| *mTurquoise2* (*Asc*I) | AAGGGTGGGCGCGGGGTGTCTAAGGGTGAGGAACTCTTC | AGCTGGGTCGGCGCGCTATTTGTAAAGCTCATCCATTCCGAG |
| *mTurquoise2-PTS1* | GCCGCCCCCTTCACCATGGTGTCTAAGGGTGAGGAACT | GGCGCGCCCACCCTTTCATAGCTTCGAAACTTTGTAAAGCTCATCCATTCCGAGG |
| pMpGE_En03  Mp*ATG13* | CTCGGTGGAGGGGTAGGGGGTCCA | AAACTGGACCCCCTACCCCTCCAC |
| pMpGE_En03  Mp*ATG14* (gRNA1) | CTCGGCAGCAGGCTGCACAGACAG | AAACCTGTCTGTGCAGCCTGCTGC |
| pMpGE_En03  Mp*ATG14* (gRNA2) | CTCGGGAAGCACGCTCCAAAGCGT | AAACACGCTTTGGAGCGTGCTTCC |
| *_pro_*Mp*MS1* | GGCCAGTGCCAAGCTTGATGAAGGGAAACAAGATGTGTCA | TTTGTACAAACTTGTTTTGAGAGATATCTCTGAACCTCGAAC |
| *_pro_*Mp*SYP2* | GGCCAGTGCCAAGCTCACGAGCGAGTGAGACACCAGAGGAG | TTTGTACAAACTTGTCCTCCTGCTTCGTGGTAAATCCTCTTC |
| *mCitrine* (*Sac*I) | GTGGTTGATAACAGCATGGTGAGCAAGGGCGAGGAGC | GATCGGGGAAATTCGTCACTTGTACAGCTCGTCCATGC |
| rbm27 (Male) | CCAAGTGCGGGCAGAATCAAGT | TTCATCGCCCGCTATCACCTTC |
| rhf73 (Female) | TGACGACGAAGATGTGGATGAC | GAAACTTGGCCGTGTGACTGA |

**Table S2. List of vectors used in LR recombination, related to STAR Methods**

S1: Ishizaki et al. (2015), S2; Minamino et al., (2021), S3; Sugano et al., (2018).

| **Expression vector** | **Entry vector** | **Ref.** | **Destination vector** | **Ref.** |
| --- | --- | --- | --- | --- |
| *_pro_*Mp*EF1α:*  *Mito*-*Citrine* | pENTR  Mito-Citrine | this study | pMpGWB303 | [S1] |
| *_pro_*Mp*SYP2:*  *Mito*-*Citrine* | pENTR  *_pro_*Mp*SYP2:Mito*-Citrine | this study | pMpGWB307 | [S1] |
| *_pro_*Mp*SYP2:*  Mp*IDH1*-*mCitrine* | pENTR MpIDH1 | this study | pMpGWB301 *_pro_*Mp*SYP2:*  Gateway-*mCitrine* | this study |
| *_pro_*Mp*DRP3:*  *3×HA-*Mp*DRP3l* | pENTR *_pro_*Mp*DRP3:3×HA-*Mp*DRP3l* | this study | pMpGWB101 | [S1] |
| *_pro_*Mp*MS1:mTurquoise2-*Mp*DRP3l^WT^* ^or^ *^K75A^* | pENTR *mTurquoise2-*Mp*DRP3l^WT^* ^or^ *^K75A^* | this study | pMpGWB101 *_pro_*Mp*MS1* | this study |
| *_pro_*Mp*ATG5:*  Mp*ATG5^res^* | pENTR *_pro_*Mp*ATG5:*Mp*ATG5^res^* | this study | pMpGWB301 | [S1] |
| *_pro_*Mp*ATG5:*  Mp*ATG5-mCitrine* | pENTR *_pro_*Mp*ATG5:*  Mp*ATG5-mCitrine* | this study | pMpGWB101 | [S1] |
| *_pro_*Mp*VAMP71:*  *mGFP*-Mp*VAMP71* | pENTR *_pro_*Mp*VAMP71:*  *mGFP*-Mp*VAMP71* | this study | pMpGWB101 | [S1] |
| *_pro_*Mp*ATG8a:*  *mTurquoise2-*Mp*ATG8a* | pENTR *_pro_*Mp*ATG8a:*  *mTurquoise2-*Mp*ATG8a* | this study | pMpGWB101 | [S1] |
| *_pro_*Mp*EF1α:*  *mTurquoise2*-*PTS1* | pENTR *mTurquoise2-PTS1* | this study | pMpGWB103 | [S1] |
| *_pro_*Mp*DUO1:*  *mTurquoise2-*Mp*GOS11* | pENTR  *mTurquoise2-*Mp*GOS11* | this study | pMpGWB101  *_pro_*Mp*DUO1* | [S2] |
| *_pro_*Mp*DUO1:*  *ST*-*mTurquoise2* | pENTR  *ST*-*mTurquoise2* | this study | pMpGWB101  *_pro_*Mp*DUO1* | [S2] |
| CRISPR/Cas9 vector | pMpGE_En03  Mp*ATG13/14* | this study | pMpGE010 | [S3] |
